# NTX250: A Modular mRNA-Based Immunotherapy Platform for HPV-Associated Cancers with Broad Applicability

**DOI:** 10.1101/2025.09.24.678202

**Authors:** Meredith L. Leong, Weiqun Liu, Nicole Fay, Diane M Da Silva, Ou Li, Chris S. Rae, Adrienne Sallets, Ruben Prins, Daniel Frimannsson, Colin McKinlay, Ray Low, Daniel J Fernandez, Gunasekaran Kannan, Pragyesh Dhungel, Sangeeta Nath, Nicole E. Peck, Elizabeth R. Webster, Ximiao Wen, Anushtha Sharma, Edward E. Lemmens, Samuel Deutsch, W Martin Kast, Ole Audun W. Haabeth

## Abstract

Infection with high-risk human papillomavirus (HPV) is a key driver of multiple HPV-associated malignancies, including cervical intraepithelial neoplasia (CIN), oropharyngeal, head and neck, and anogenital cancers. Despite the high efficacy of prophylactic HPV vaccines, a substantial population remains at ongoing risk due to issues related to vaccine accessibility, awareness, or personal choice. Current treatments, such as the loop electrosurgical excision procedure (LEEP), address lesions but do not eliminate persistent HPV infection and are associated with potential complications. Moreover, individuals treated for HPV-related disease are at increased risk for subsequent HPV-associated malignancies. These factors underscore an urgent need for effective, non-invasive therapeutic strategies capable of targeting and eradicating persistent HPV infections.

Here we introduce NTX250, an innovative mRNA-based therapeutic platform designed to deliver HPV type 16 (HPV16) E6-E7 antigens, combined with immunomodulators human interleukin-12 (IL-12) p70 and engineered human LIGHT (LIGHT). The mRNAs are co-delivered via lipid and ionizable peptoid nanoparticles referred to as Nutshell™ formulation for intralesional administration.

Our preclinical evaluation demonstrates that localized delivery of NTX250 in HPV16-transformed tumor-bearing models results in complete tumor regression and the induction of durable, antigen-specific immune memory. Moreover, this modular mRNA nanoparticle approach combining tumor-specific antigens and immunomodulators is adaptable beyond HPV, with potential applications targeting neoepitopes or other tumor-associated antigens, thus offering a versatile platform for immunotherapy across a broad range of indications. These findings highlight the potential of this strategy as a broadly applicable, non-surgical immunotherapeutic capable of inducing robust and specific anti-tumor responses.

## Introduction

mRNA therapeutic modalities facilitate the transient and safe delivery of messenger RNA encoding therapeutic proteins that are expressed directly in patient cells. Recent clinical findings affirm the effectiveness of mRNA nanoparticles in eliciting robust and sustained antigen-specific immune responses to mRNAs encoding an antigenic protein or peptide, coupled with an immunogenic adjuvant effect stemming from the drug product (comprising both mRNA and nanoparticles)^1^. However, the nature of this adjuvant effect remains largely arbitrary and not necessarily targeted toward specific immune responses.

Development efforts for nanoparticles have predominantly concentrated on the safe and efficient delivery of nucleic acid cargo, prioritizing factors such as stability, delivery efficiency, and tolerability. The recent successes of SARS-CoV-2 mRNA vaccines, along with the generation of neoantigen-specific T cell responses in patients treated with personalized neoantigen mRNA nanoparticle vaccines, underscore the potential of mRNA nanoparticle therapeutics to induce both B and T cell antigen-specific immune responses^2–4^. However, these mRNA nanoparticle products often lack a mechanism to intentionally guide immune responses toward the specific responses historically associated with improved clinical outcomes in cancer patients.

Tumors are complex multicellular structures capable of attenuating and redirecting immune responses to evade anti-tumor activity. While generating tumor-specific immune responses is a substantial achievement, it is, in most cases, insufficient to attain durable therapeutic outcomes (reviewed^5,6^). Indeed, most personalized cancer vaccines necessitate the concurrent application of additional treatments such as checkpoint inhibitors to overcome tolerance/suppression and effectively reduce tumor burden^7^.

The cargo and localization of expression are critical components that can empower mRNA nanoparticle drugs to surmount these limitations. Intracellular delivery and expression of proteins encoded by mRNA present an unique opportunity to strategically direct various elements of the immune response to specific tissue and cellular compartments that are more conducive to drive therapeutic activity. Proteins synthesized from mRNA can be targeted toward intracellular locations, integrated into cellular membranes, or secreted into the local microenvironment. By harnessing these distinctive properties, drug designers can optimize the expression of proteins to their most effective compartments, thereby fostering synergistic immune responses. Additionally, mRNA nanoparticle therapeutics allow for wider combinations of agents, such as pairing antigens with immunomodulators, to enhance and tailor immune responses more precisely. This approach necessitates a delivery strategy where expression is primarily localized in the site of injection without significant “leakage” to systemic organs which has been observed for many existing lipid nanoparticles.

Tumors evolve mechanisms to ensure their nutrient acquisition, oxygenation, avoidance of immune elimination, and restriction of tissue growth^8^. The continuous advancement of immunotherapies hinges on the ability of such agents to overcome the immune suppression and exclusion orchestrated by the tumor microenvironment (TME). A strategic approach to modifying the TME and generating positive feedback loops that enhance immune support and recruit anti-tumor effector immune cells is through the localized administration of immunotherapeutic agents. The historical context of intratumoral immunotherapy administration reflects a paradox of pre-clinical successes contrasted by clinical failures^9^. Various factors contribute to the lack of consistent clinical success, including the absence of suitable models that accurately mimic the evolution of human tumors, the misconception that targeting a single aspect of tumor biology or a singular arm of the immune response will suffice for durable efficacy, a misunderstanding of local versus systemic drug exposure, lack of standardized injection protocols, and the difficulty in striking a balance between effective immune stimulation and harmful overstimulation.

In this study, we present NTX250, a locally administered, carefully engineered multimodal mRNA-based immunomodulatory vaccine that demonstrates the capacity for the complete and sustained clearance of large, established, clinically relevant, and difficult-to-treat tumors in murine models. Additionally, we show that NTX250 elicits robust antigen-specific T cell responses in cynomolgus macaques and enhances in vitro reactivation and proliferation of human HPV16-specific CD8 T cells derived from patients with cervical intraepithelial neoplasia (CIN).

The delivery of NTX250 is facilitated by a proprietary nanoparticle system with good tolerability and in vivo transfection efficacy, which, in contrast to many lipid nanoparticles, remains preferentially localized to drive expression at the site of injection^10^. This multimodal mRNA formulation comprises an enhanced chimeric HPV16 E6 and E7 antigen, a secreted single-chain interleukin-12 (IL-12) p70, and a membrane-anchored Tumor Necrosis Factor Superfamily Member 14 (TNFSF14, also known as LIGHT). These three mRNAs are co-formulated in a 1:1:1 ratio (w/w) in a Nutshell™ nanoparticle^11^ optimized for intra-lesional administration referred to as DVI-140, facilitating robust local transfection while minimizing secondary leakage to organs in circulation such as the liver. The chimeric E6-E7 protein is engineered to enhance antigen presentation by localizing to late endosomes/MHC class II sorting complexes, achieved through the fusion of the E6-E7 protein with a human CD1d secretion signal at the N-terminus and the transmembrane domain and short cytoplasmic tail of CD1d at the C-terminus. The integrity of IL-12p70 is guaranteed through the construction of a single-chain IL-12p40-IL-12p35 molecule connected by a flexible G4S linker. The potency of LIGHT is amplified through the removal of the enzymatic cleavage site responsible for its membrane release, replaced with a rigid linker to maintain functionality.

Overall, NTX250 was meticulously designed to induce robust HPV16-specific immune responses, promote immune cell infiltration within injected tissues, and remodel the TME from an immunosuppressive to an immuno-enhancing profile. This novel genetic drug offers a potential therapeutic strategy for individuals with active HPV infections to clear lesions and resolve the underlying infection across a spectrum of HPV-related precancerous conditions and cancers.

## Results

### Enhanced Antigen Presentation and T Cell Responses Through Subcellular Targeting of HPV16 E6-E7 Fusion Antigens

HPV16 E6 and E7 are oncogenic proteins that play a key role in the development of cervical cancer and other HPV-associated malignancies. E6 promotes the degradation of p53, a crucial tumor suppressor protein, leading to impaired apoptosis and uncontrolled cell growth^12^. It also activates telomerase, contributing to cellular immortality^13,14^. E7 binds to and inactivates retinoblastoma protein (pRb), releasing E2F transcription factors that drive unregulated cell cycle progression^13^. Together, E6 and E7 disrupt normal cell cycle control, allowing HPV-infected cells to evade growth suppression and potentially become cancerous and identifying them as critical targets for therapeutics. We designed a mutated fusion protein of the HPV16 E6 and E7 that retains the immunodominant epitopes but abrogated the oncogenic activity^15^, and we evaluated several antigen designs to enhance MHC class I presentation and to elicit HPV16 E6-E7 specific CD8+ T cell responses.

#### Optimization of HPV16 E6-E7-Specific T Cell Responses

Given the well-characterized immunodominant E7_49-57_ epitope in C57BL/6 mice (RAHYNIVTF)^16^, we focused on generating robust CD8+ T cell responses in murine models. To enhance CD8+ effector T cell induction and persistence, a process reliant on CD4+ T cell help, we incorporated the universal Pan HLA-DR T helper epitope (PADRE) into our E6-E7 antigen design^17^. Mice received two intramuscular (IM) injections 14 days apart of 2µg of Nutshell^TM^ DVI-140 formulated mRNA encoding fusion molecule of HPV16 E6-E7 (E6-E7) or a design that also included PADRE which has a high affinity for human and mouse MHC II molecules (PADRE-E6-E7). Seven days post-boost, antigen-specific CD8+ and CD4+ T cell responses were assessed. Inclusion of the PADRE significantly enhanced E7-specific CD8+ responses (Supplementary Figure 1A) and elicited PADRE-specific CD4+ T cell responses (Supplementary Figure 1B). These findings support previous observations that CD4+ T-helper epitopes aid CD8+ T cell priming through dendritic cell (DC) licensing and cytokine support^18,19^.

#### Subcellular Targeting to Enhance Antigen Presentation

We next examined antigen localization strategies to improve antigen processing and MHC presentation. Constructs encoding a fusion protein of E6-E7 were designed for cytosolic expression with no signal peptide (E6E7), for endoplasmic reticulum (ER) retention (E6E7-KDEL), or for endolysosomal trafficking via trafficking domains such as MITD^20^ (E6E7-MITD) and CD1d (E6E7-hCD1d). Cytosolic and ER-retained constructs were expected to favor MHC class I presentation^21^, while endolysosomal-targeted constructs aimed to enhance both MHC class I and class II loading.

To assess in vitro antigen presentation of the different constructs, we employed an HLA-A2-restricted E7_11-21_-specific TCR-transgenic (TCR-Tg) Jurkat reporter cell line to evaluate antigen presentation. Jurkat cells expressing Lucia under an NFAT promoter secreted Lucia upon recognition of E7_11-21_ presented on HLA-A2 molecules. HLA-A2+ HeLa, monocyte-derived DCs, and THP-1 cells were transfected with E6-E7 constructs and co-cultured with TCR-Tg Jurkat cells. Lucia expression was measured at 48-hours post-transfection. Endolysosomal trafficking domains, MITD and hCD1d, significantly improved antigen presentation in all cells tested compared to cytosolic and ER-retention strategies (Supplementary Figure 1C).

To validate findings in primary human cells, we designed similar constructs encoding HLA-A2 epitope CMV pp65_495-503_ (NLVPMVATV). Peripheral blood mononuclear cells (PBMCs) from two HLA-A2+ CMV-reactive donors were transfected with 100 ng CMVpp65, CMVpp65-MITD, or CMVpp65-hCD1d mRNA formulated in Nutshell™ nanoparticles. IFN-γ expression in CD4+ and CD8+ T cells was measured 12 hours post-transfection. The highest IFN-γ responses were observed in PBMCs treated with CMVpp65-hCD1d mRNA (Supplementary Figure 1D).

To assess construct immunogenicity in vivo, mice received two IM injections of 2µg E6E7, E6E7-KDEL, E6E7-MITD, or E6E7-hCD1d mRNA formulated in Nutshell™ nanoparticles. Seven days post-boost, HPV16 E7-specific CD8+ T cell responses were assessed by ELISpot and tetramer staining. MITD significantly enhanced E7-specific CD8+ responses compared to other constructs (Supplementary Figure 1E). In contrast, human CD1d (E6E7-hCD1d) did not enhance responses, likely due to species incompatibility of the human CD1d sequence in mice. To address this, a murine CD1d (mCD1d) construct was generated (E6E7-mCD1d) and demonstrated comparable efficacy to MITD in vivo (Supplementary Figure 1F).

#### Subcellular Localization of HPV16 E6-E7-CD1d Proteins

To confirm the activity of the endolysosomal trafficking domain CD1d, we conducted subcellular localization experiments. HeLa cells were transfected with mRNA encoding different E6-E7 constructs, and expression of the E6-E7 fusion protein was assessed by flow cytometry and confocal microscopy using an anti-E7 antibody. Flow cytometry analysis revealed that only MITD-containing E6-E7 proteins localized to both the cell surface and intracellular compartments, whereas cytosolic, ER-retained, and CD1d-containing proteins were detected only intracellularly (Supplementary Figure 2A).

Further evaluation using confocal microscopy demonstrated that both MITD- and CD1d-containing E6-E7 proteins accumulated in the multivesicular bodies of late endosomes (Supplementary Figure 2B). These findings confirm that CD1d directs antigen expression into compartments that facilitate both MHC class I and MHC class II presentation. As most MHC class II-presented epitopes originate from extracellular antigens acquired via phagocytosis or endocytosis, our results suggest that including CD1d in antigen designs provides an effective strategy to achieve MHC class II presentation of intracellular antigens.

The combination of in vitro and in vivo results suggests that the CD1d endolysosomal trafficking domain significantly enhances mRNA-encoded antigen performance as evidenced by enhanced TCR-specific signaling (Supplementary Figure 1C) and the increased generation of antigen-specific T-cells in vivo relative to constructs without trafficking domains or ER retention sequences (Supplementary Figure 1E). Confocal microscopy data confirms that the addition of endolysosomal domains leads to antigen localization in late endosomes, a subcellular compartment where MHC-I and II loading takes place. While CD1d outperformed MITD in human cells, in vivo murine studies showed equivalent efficacy between mCD1d and MITD, underscoring the importance of species-specific compatibility in antigen trafficking strategies.

#### IL-12 and LIGHT synergize to enhance T cell activity and expansion

In the design of NTX250, we aimed to evaluate the potential synergistic activity of IL-12 and LIGHT, leveraging IL-12’s multifaceted immunostimulatory capabilities as a pluripotent cytokine and LIGHT’s role as a potent T cell co-stimulator, to enhance T cell activation and bolster anti-tumor immune responses. In vitro, we wanted to assess the direct effect of IL-12 and LIGHT, both independently and in combination, on T cell activation and expansion. IL-12p70 is a heterodimeric pro-inflammatory cytokine composed of p35 and p40 subunits, primarily produced by activated antigen-presenting cells, such as dendritic cells and macrophages. IL-12 plays a pivotal role in the regulation of innate and adaptive immunity by promoting the differentiation of naïve CD4+ T cells into Th1 effector cells and enhancing the cytotoxic function of CD8+ T cells. It stimulates T cells and natural killer (NK) cells to produce interferon-gamma (IFN-γ), which further amplifies the Th1 immune response and supports anti-tumor activity^22–24^. IL-12 also enhances the proliferation and survival of activated T cells, making it a critical immunomodulatory cytokine in cancer immunotherapy. Its potent immunostimulatory properties have led to its incorporation in various therapeutic platforms aimed at reprogramming the tumor microenvironment toward a pro-inflammatory, immune-active state. LIGHT, a member of the tumor necrosis factor (TNF) superfamily, is a potent immunostimulatory molecule that plays a key role in T cell co-activation, tumor immunity, and the modulation of the tumor microenvironment. LIGHT interacts primarily with two receptors: HVEM (herpesvirus entry mediator) and LTβR (lymphotoxin β receptor).

Through these interactions, LIGHT enhances T cell proliferation, survival, and effector function, while also promoting dendritic cell maturation and cytokine production^25,26^. In tumor models, LIGHT expression has been shown to increase CD8+ T cell infiltration, promote a pro-inflammatory microenvironment, and inhibit tumor growth by enhancing cytotoxic T cell activity^25^. LIGHT also induces chemokines like CXCL9 and CXCL10, supporting T cell trafficking to tumor sites, making it a valuable component in cancer immunotherapy strategies.

To further characterize the interaction between IL-12 and LIGHT, we studied the effect of IL-12, LIGHT or both in human primary T cells isolated from eight different healthy donors. CellTrace^TM^ Violet-labeled primary human CD3+ T cells were co-cultured with either untransfected or mRNA-transfected HEK293 cells expressing non-coding mRNA, IL-12, LIGHT, or both IL-12 and LIGHT. Agonistic CD3 antibody was included to mimic TCR engagement and provide signal 1. After 4 days of co-culture, T cell proliferation was assessed via flow cytometry. The percentage increase in proliferated CD8+ T cells relative to CD3-only controls was calculated for each donor. In five of the eight donors, T cells exposed to both LIGHT and IL-12 showed increased activation—as indicated by CD25 expression, higher numbers of cells proliferating, and greater numbers of division cycles (Figure 1A-B, Supplementary Figure 3A-B). This demonstrated that the combination of IL-12 and LIGHT significantly enhanced T cell expansion and survival.

**Figure 1:**
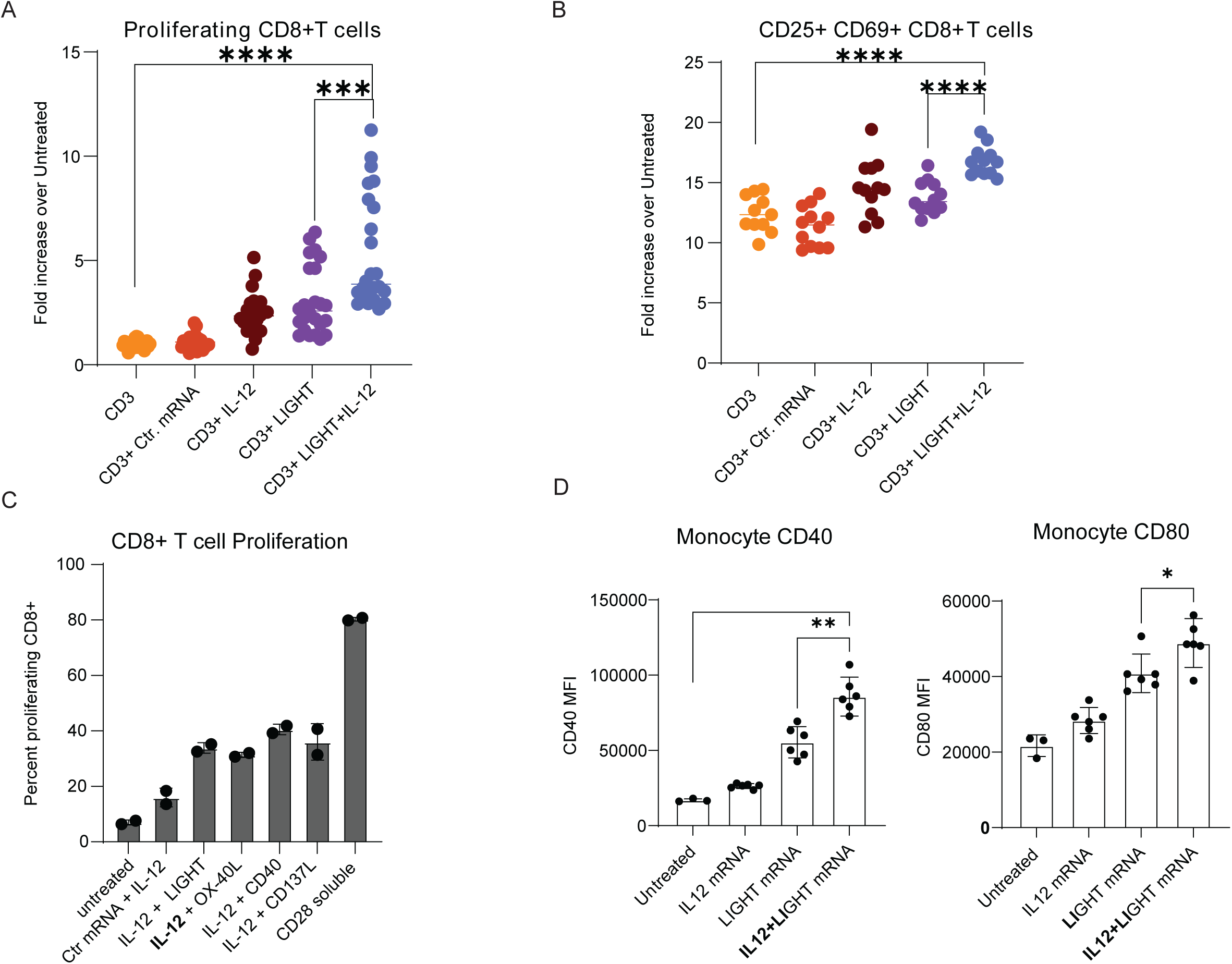
IL-12 and LIGHT synergize to enhance immune activation. **A, B.** HEK293 cells were transfected with mRNA encoding IL-12, LIGHT, both (LIGHT+IL-12), or a non-coding mRNA (Ctr mRNA). Purified CD3+ T cells from healthy donors (n=8) were incubated with anti-CD3 antibody (CD3+) and co-cultured with transfected HEK293 cells. Each condition was performed in triplicate, with data pooled across donors. After 4 days of incubation, proliferating CD8+ T cells were assessed by CellTrace Violet (CTV) dilution **(A)** and number of CD25+ CD69+ CD8+ T cells quantified **(B)** by flow cytometry. Fold increase over untreated was calculated for each donor. **C.** HEK293 cells were transfected with mRNA encoding IL-12 in combination with mRNAs encoding LIGHT, OX-40L, CD40, or CD137L, or with control mRNA (Ctr mRNA). Purified CD3+ T cells from two donors were incubated with anti-CD3 and co-cultured with the transfected HEK293 cells. T cell proliferation was measured via dye dilution after 4 days. Incubation with soluble agonistic CD28 (CD28 soluble) served as a positive control. **D.** From 5 donors, CD14+ monocytes were isolated and treated with mRNA encoding IL-12, LIGHT, or both mRNA (IL12+LIGHT) or lef t untreated. After 48 hours, surface expression levels of CD40 **(left)** and CD80 **(right)** on monocytes were measured by flow cytometry. Error bars represent mean ± SD, with each symbol indicating the mean value from technical replicates for individual donors. Statistical significance was determined by two-tailed, Student’s t-test: *\*p<0.05, **p<0.01, ***p<0.001, ****p<0.0001*

To evaluate the co-stimulatory capacity of LIGHT relative to other molecules, mRNAs encoding OX-40L, CD137/4-1BB, and CD40 were tested in combination with IL-12 mRNA and soluble aCD3 in the same co-culture assay. On day 6, CD8+ T cell proliferation was measured. LIGHT + IL-12 co-stimulation was comparable to that of OX-40L and 4-1BB, while CD40 + IL-12 yielded a slightly higher response. Soluble agonistic aCD28 served as a positive control and induced the highest proliferation (Figure 1C).

Additionally, IL-12 and LIGHT synergistically activated CD14+ monocytes from five donors. Following 48 hours of co-culture with mRNA-transfected HEK293 cells, monocytes showed significantly increased surface expression of CD40 and CD80, indicating enhanced activation and maturation (Figure 1D).

The observed synergistic effects of IL-12 and LIGHT in promoting T cell activation, proliferation, and monocyte maturation support their strategic inclusion in NTX250, aiming to maximize immune engagement and therapeutic efficacy through combined immunostimulatory mechanisms.

### Intratumoral Delivery of mNTX250 Induces Complete Tumor Regression

mNTX250 is a murine version of NTX250 that includes species-specific mRNAs encoding E6-E7- mCD1d, murine IL12p70 and murine LIGHT encapsulated in Nutshell™ nanoparticles. We evaluated the efficacy of mNTX250 in a preclinical tumor model, C3.43, a progressive subclone derived from the HPV16-transformed C3 B6 mouse embryo cell line, which expresses HPV16 E6 and E7 under viral regulation at levels similar to those observed in HPV infected tissues in patients^27–30^. C57BL/6 mice were inoculated with subcutaneous tumors 15 to 18 days before treatment initiation. Mice received three intratumoral (IT) or intramuscular (IM) injections of mNTX250 (1.5µg total mRNA) at seven-day intervals. Control animals received 1.5µg of non-coding mRNA.

Both IT and IM administration led to a significant induction of HPV16 E7-specific CD8+ T cells compared to control-treated mice (Figure 2A). However, IT administration induced significantly higher E7-specific CD8+ T cell responses than IM administration and resulted in complete long-term tumor eradication, leading to significantly improved overall survival (Figure 2A-C). These findings underscore the superior efficacy of IT administration in generating potent anti-tumor immunity and achieving durable tumor regression.

**Figure 2:**
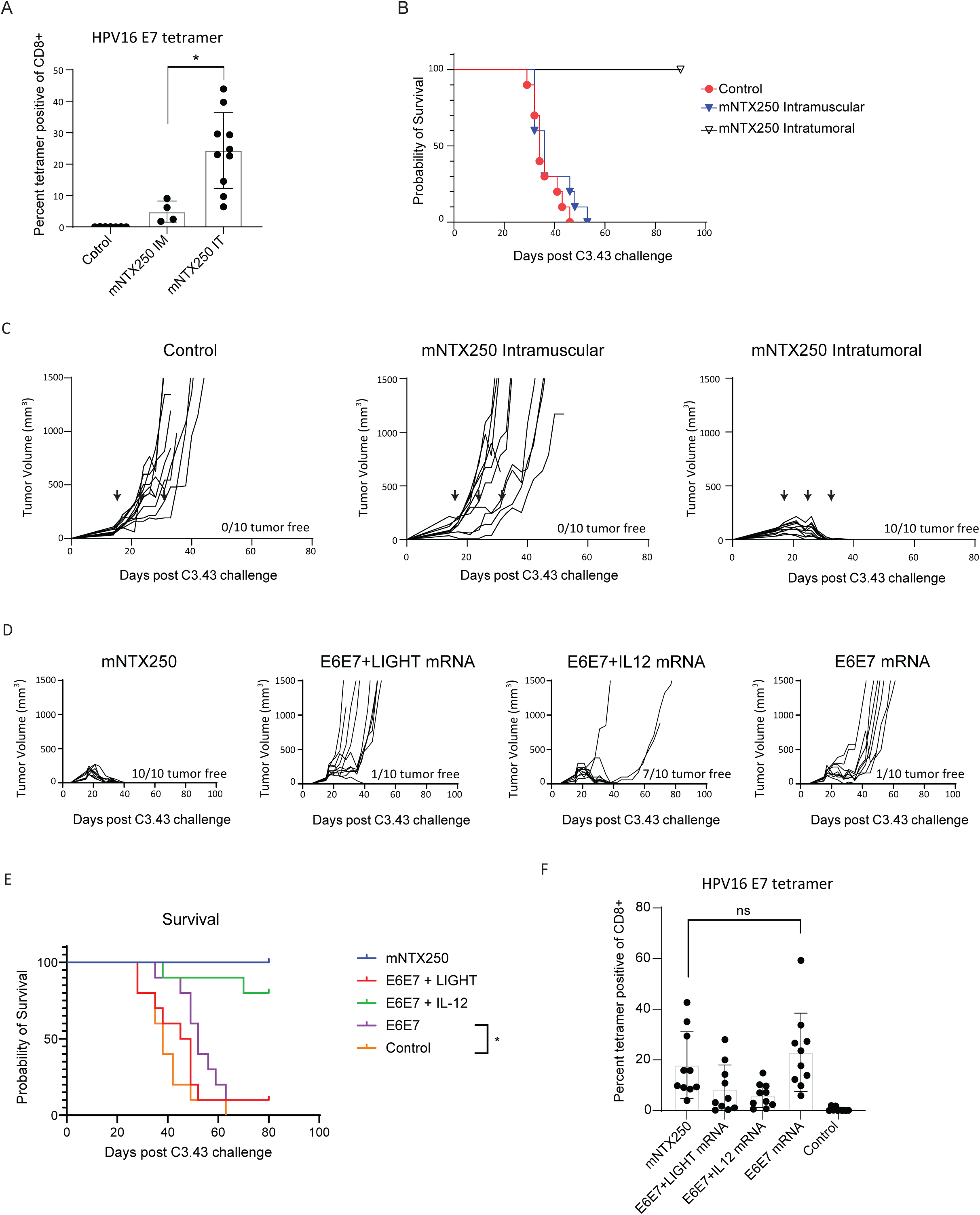
CD8+ T-cell dependent tumor efficacy following intratumoral delivery of mNTX250. C57BL/6 mice (n=10) were implanted subcutaneously with C3.43 tumor cells. Once tumors reached a volume of 50–100 mm³, treatments began and were administered once weekly for three doses (indicated by arrows). **A, B, C.** Mice received 1.5µg mNTX250 intramuscular or intratumoral or treatment control. **D, E, F.** Mice received 1.5µg of mNTX250 or Nutshell™ formulations of mRNAs encoding HPV16 E6-E7 alone (E6E7 mRNA) or in combination with mRNA encoding LIGHT (E6E7+LIGHT mRNA) or IL-12 (E6E7+IL12 mRNA). Seven days after the final dose, PBMCs were collected, and HPV16 E7_49–57_-specific responses were quantified as percent of CD8+ by tetramer analysis **(A, F)**. **A.** Mice were monitored daily for survival **(B, E)** and tumor growth was measured over time. Number of tumor-free animals is indicated on each graph **(C, D)**. Untreated C3.43 tumor-bearing animals were used as controls for tumor growth and survival **(B, C, D, E)**. **A**. Control group for tetramer consists of 3 tumor-bearing controls and 4 naïve C57BL/6 mice. Error bars represent mean ± SEM. Statistical significance was determined by two-tailed, Student’s t-test: \**p<0.05, ns: not significant*.

#### IL-12 and LIGHT Are Required for NTX250 Efficacy

To determine the requirement for IL-12p70 and LIGHT mRNA in the mNTX250 formulation, we treated mice with large, established C3.43 tumors with three intratumoral doses of Nutshell™ nanoparticles containing HPV16 E6-E7 mRNA alone, HPV16 E6-E7 + IL-12 mRNA, HPV16 E6-E7 + LIGHT mRNA, or the mNTX250 full triple-mRNA formulation. When applicable, mRNAs were formulated at 1:1 or 1:1:1 ratios, ensuring the total mRNA dose remained constant at 1.5µg.

Consistent with previous results, three doses of mNTX250 resulted in rapid and complete tumor clearance. In contrast, HPV16 E6-E7 mRNA alone did not achieve tumor clearance, nor did the combination of LIGHT and HPV16 E6-E7 mRNA. The addition of IL-12 to HPV16 E6-E7 mRNA led to complete, long-term tumor clearance in 70% of treated mice (Figure 2D-E). Notably, all treatment groups exhibited a significant increase in circulating HPV16 E7-specific CD8+ T cells compared to untreated controls (Figure 2F). These findings suggest that modulation of the tumor microenvironment is critical for achieving robust therapeutic responses.

#### mNTX250 Induces CD8 T Cell-Dependent Long-Term Anti-Tumor Immunity

To assess the durability of mNTX250-induced immunity, the 10 surviving mice from the mNTX250 IT-treated group were divided into two cohorts 89 days after the initial inoculation. Five animals received CD8+ T cell depletion via repeated dosing with anti-CD8 antibodies, while the other five received isotype control antibodies. Complete CD8+ depletion was verified by flow cytometric analysis of circulating CD8+ T cells (Supplementary Figure 4A). Once depletion was confirmed, mice were rechallenged with C3.43 tumor cells. As tumor growth control, five naïve animals were also challenged with C3.43 tumors. All CD8+ T cell-depleted and treatment-naïve mice developed tumors, whereas all isotype control-treated mice rejected the rechallenge (Supplementary Figure 4B). These results confirm that mNTX250 induces long-term protective immunity against HPV16-driven tumors, which is critically dependent on CD8+ T cells.

### Two Doses of mNTX250 Are Sufficient for Tumor Eradication and Long-Term Protection

To further evaluate mNTX250 efficacy, C57BL/6 mice with established subcutaneous C3.43 tumors received two intratumoral injections of mNTX250 at three dosage levels (1.5µg, 0.45µg, and 0.045µg), administered seven days apart. Control groups received either non-coding mRNA or mNTX151, Nutshell™ nanoparticles encapsulating only IL-12p70 and LIGHT mRNA. Tumor growth and survival were monitored biweekly.

All dose levels of mNTX250 resulted in complete long-term tumor eradication in 90 to 100 percent of the treated animals (Figure 3A-B). Administration of mNTX151 was only able to eradicate tumors at the 1.5µg and the 0.45µg. Treatment with 1.5µg and 0.45µg of mNTX250 induced robust dose dependent HPV16 E7-specific CD8+ T cell responses, as indicated by circulating tetramer-positive CD8+ T cells, whereas NTX-151 treatment did not significantly enhance E7-specific CD8+ responses (Figure 3C). In a follow-up study, surviving mNTX250-treated animals rejected a subsequent challenge with syngeneic HPV16-expressing TC-1 tumors, while treatment-naïve and mNTX-151-cured animals developed tumors (Figure 3D). These results demonstrate that mNTX250 not only eliminates large established tumors but also provides durable memory protection against HPV16-associated malignancies. Notably, local treatment with the two immunomodulators alone in mNTX151 was able to clear tumors at the higher doses but could not prevent growth of tumors by rechallenge with the HPV16 E6-E7 expressing TC-1 cell line.

**Figure 3:**
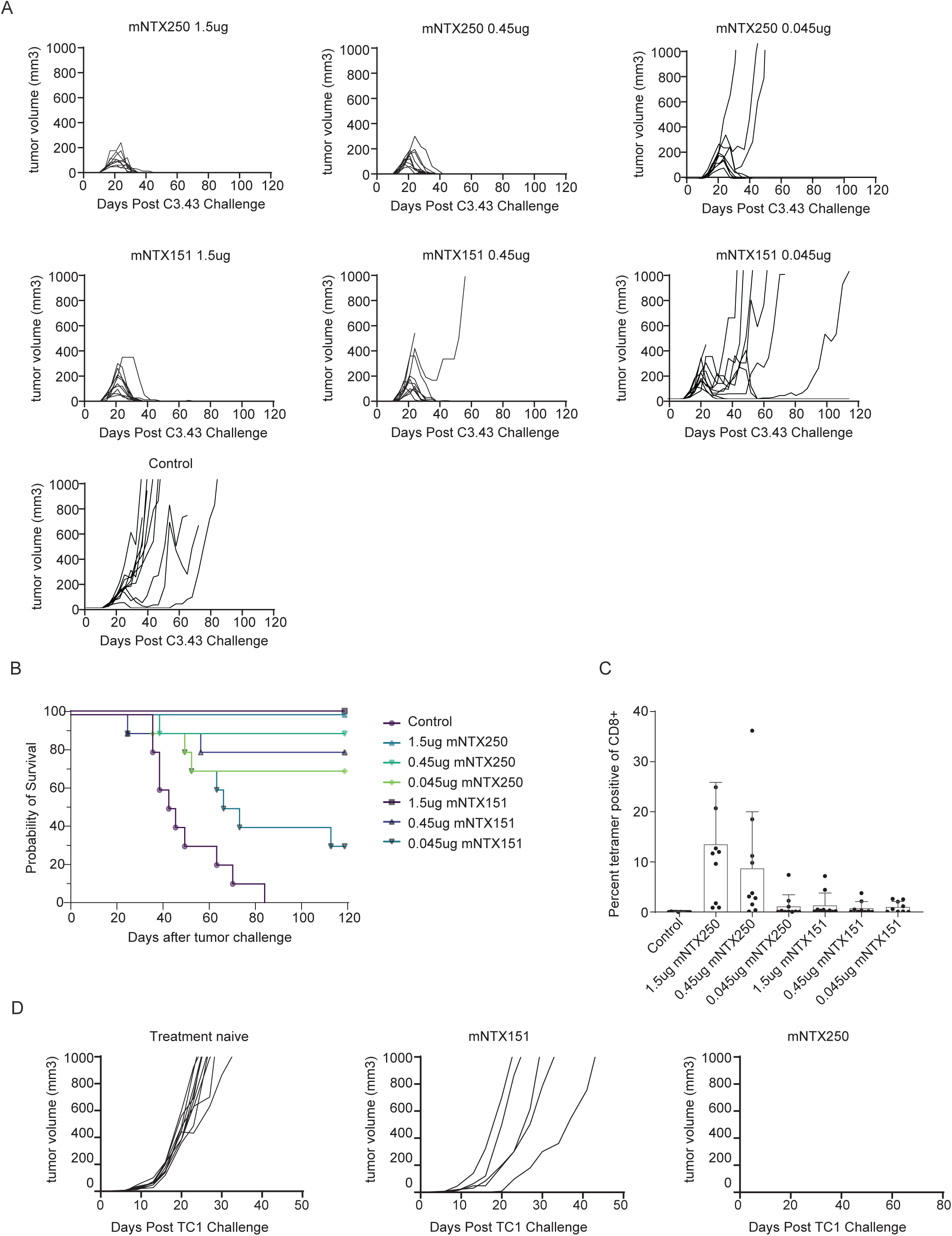
Robust efficacy across a dose range of mNTX250. C57BL/6 mice were implanted subcutaneously with C3.43 tumors. Starting when tumor volumes were between 50-100mm^3^, animals (n=10/group) were administered intratumorally once weekly for two doses with 1.5, 0.45, 0.045µg of mNTX250 **(A, upper panels)** or mNTX151 which is a Nutshell^TM^ formulation of mRNAs encoding mIL12 and mLIGHT **(A, middle panels)**, or control. Tumor growth was measured (**A)** and survival monitored over time **(B).** Four days after second dose, HPV16 E7_49-57_-specific responses were measured in PBMCs by tetramer analysis **(C)**. **D.** Tumor-free animals previously treated with mNTX250 (n=15; right panel) or with mNTX151 (n=11; middle panel) were rechallenged subcutaneously with TC-1 tumor cells and tumor growth measured. For rechallenge control group, naïve C57BL/6 (n=10, Treatment naïve; left panel) were implanted with TC1 and monitored over time. Error bars represent the mean ± SEM.

### Immunogenicity, Tumor Microenvironment Modulation, and Immune Profiling of mNTX250

To evaluate the immunogenic effects of mNTX250, C57BL/6 mice with established subcutaneous C3.43 tumors received two doses of mNTX250 at 4-day intervals with dose levels ranging from 1.5 to 0.045 µg of drug product. Animals were euthanized one day after the second dose for mechanistic studies and immune response profiling. Tumors were excised and processed for tumor-infiltrating lymphocyte (TIL) isolation and tumor supernatant collection for cytokine analysis and flow cytometry. The immunogenicity of mNTX250 was further assessed via flow cytometry of PBMCs, and the frequencies of HPV16-E7-specific CD8+ T cells were quantified across all treatment groups. A dose-dependent increase in circulating HPV16-E7-specific CD8+ T cells was observed (Figure 4A), along with enhanced tumor infiltration of CD8+ and CD4+ T cells and natural killer (NK) cells. (Figure 4B). The cytokine profile shows a Th1-skewed immune response, characterized mainly by increased secretion of IFNγ in a dose-dependent manner compared to untreated controls (Figure 4C). Notably, a marked reduction in FoxP3-positive CD4+ regulatory T cells (Tregs) was observed in all mNTX250-treated animals relative to untreated controls (Figure 4D), indicating a favorable immunomodulatory shift toward an anti-tumor immune microenvironment. These findings clearly demonstrate the ability of mNTX250 to modulate tumor microenvironment and enhance immune responses for overall therapeutic efficacy.

**Figure 4:**
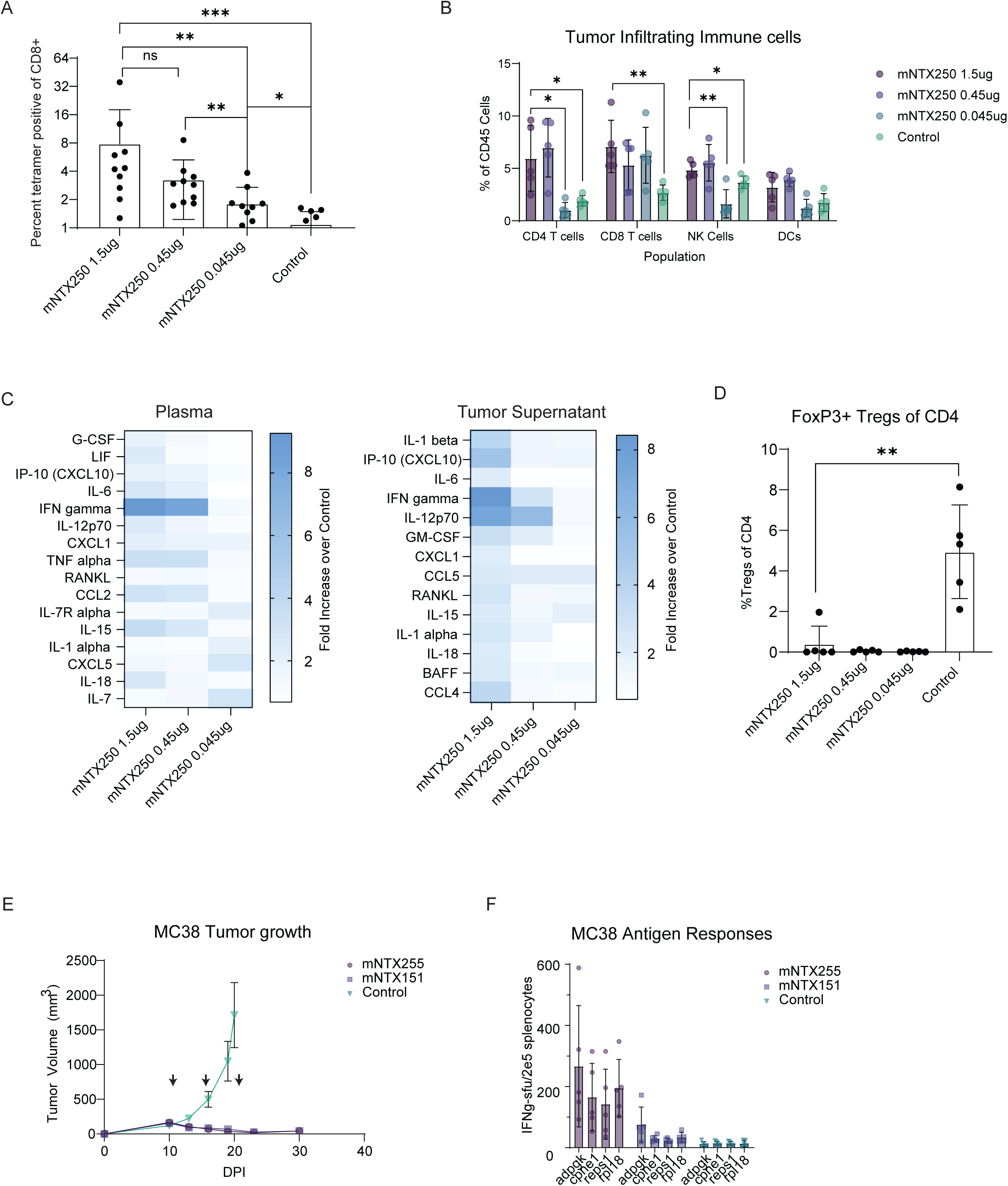
mNTX250 favorably modulates tumor microenvironment. **A, B, C, D.** C57BL/6 mice (n=10) were implanted subcutaneously with C3.43 tumors. 1.5, 0.45, 0.045µg of mNTX250 was administered intratumorally once weekly for two doses. Untreated tumor-bearing animals (n=10) were used as controls. One day after the second dose, tumor and blood were collected and processed to tumor-infiltrating cells, tumor supernatant, plasma, and PBMCs. HPV16 E7_49-57_-specific responses as percent of CD8+ T cells were measured in PBMCs by tetramer analysis **(A)**. Infiltrating CD4+, CD8+, NK and DCs were measured in tumors by flow cytometry **(B)**. Cytokines were measured in plasma **(left)** and tumor supernatants **(right)** using multiplex assay and analyzed as fold increase of treatment mean over vehicle control mean **(C)**. Tumor-infiltrating FoxP3+ Treg cells were assessed as percent CD4+ T cells in tumors by flow cytometry **(D)**. **E, F.** C57BL/6 mice (n=10) were implanted subcutaneously with MC38 tumors. Mice were administered 5µg of mNTX255 (Nutshell^TM^ formulation of mRNAs encoding MC38 neoepitopes, mIL-12, and mLIGHT), mNTX151 (Nutshell^TM^ formulation of mRNAs encoding mIL-12 and mLIGHT), or non-coding mRNA control IT once weekly for three doses and tumor growth monitored over time **(E)**. Seven days after third dose, immune responses to MC38 neoepitopes adpgk, cpne1, reps1, and rpl18 were measured in spleens as spot-forming units (SFU) by IFNγ ELISpot. Spleens from control animals were collected at time of euthanization, processed to single cells, and frozen until samples were collected from remaining animals. **(F)**. Error bars represent the mean ± SEM. p values were determined by two-tailed, student’s t-test. *\*p < 0.05, **p < 0.01, ***p < 0.001*.

### Generalizing the approach to non-HPV driven solid tumors

To evaluate the therapeutic potential of the NTX250 platform to other solid tumors, we replaced the HPV E6-E7-CD1d antigen with an antigenic construct comprising MC38 specific neoepitopes^31^ to produce mNTX255 (formulation of mRNAs encoding mIL-12, mLIGHT, and MC38 neoepitopes). Animals with established subcutaneous MC38 tumors were administered mNTX255, mNTX151 (formulation of mIL-12 and mLIGHT) or a control formulation three times once a week intratumorally and tumor growth monitored. Seven days after last dose, mNTX255 and mNTX151 animals were euthanized, and neoepitope-specific immune responses assessed.

Strong tumor growth suppression was observed in both the mNTX255 and mNTX151 treatment groups compared to control treated, with tumor control levels being similar between these groups. This indicates that these immunomodulators have broad applicability across different tumor types (Figure 4E). Importantly, analysis of splenocytes demonstrated a robust induction of MC38 neoepitope-specific T cell responses, as measured by IFNγ release assays. The generation of these specific responses was dependent on the inclusion of neoepitope-encoding mRNA, underscoring the necessity of neoepitope delivery in eliciting targeted anti-tumor immunity (Figure 4F). These results support the potential of the mNTX250 platform for personalized immunotherapy, capable of inducing potent neoepitope-specific T cell responses crucial for effective tumor control.

### NTX250 Induces Robust HPV cellular immunogenicity in Non-Human Primates and CIN Patient-Derived PBMCs

To evaluate the immunogenicity of NTX250 in vivo, we first assessed its activity in non-human primates (NHPs). Cynomolgus macaques were immunized three times intramuscularly weekly with either a low (375 µg) or high (1125 µg) dose of NTX250 (n=2 per group). Blood samples were collected seven days following the final dose, and HPV16 E6-E7-specific T cell responses were quantified by IFNγ ELISpot. Both dose groups demonstrated robust induction of E6-E7-specific T cell responses (Figure 5A). No appreciable dose-dependent differences were observed. No adverse local or systemic effects were observed in any of the animals, but detailed histological studies were not performed at this stage.

**Figure 5:**
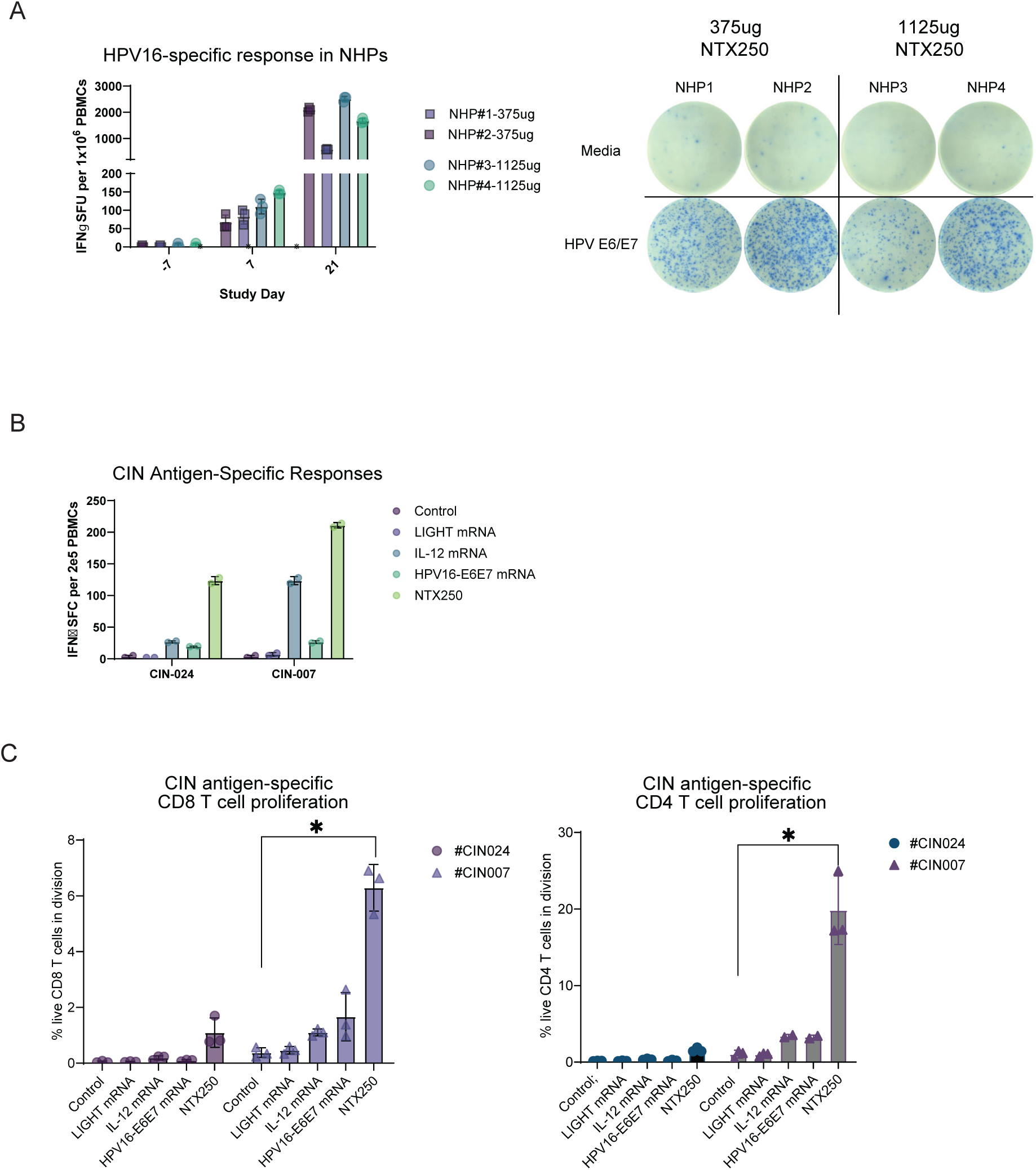
Activation of T cells by NTX250 in NHPs and with human PBMCs. **(A)** On day 0, cynomolgus macaques (n=2/group) were administered 375ug (NHP#1 and #2) or 1125ug (NHP#3 and #4) of NTX250 intramuscularly three doses separated by 21 days. On days -7, 7, and 21, PBMCs were collected, and HPV16-specific immune responses were measured as IFNγ spot-forming units (SFU) by ELISpot using an overlapping HPV16 E6-E7 peptide library (HPV E6-E7) and media only as a negative control (media). Representative images from one well per stimulated sample in IFNγ ELISpot are shown. **B, C.** PBMCs from two CIN donors (#CIN024, #CIN007) were treated with NTX250 or Nutshell^TM^ formulations of mRNA encoding LIGHT, IL12, or HPV16 E6-E7 or a non-coding mRNA control. Cells were incubated on anti-IFNγ coated ELISpot plate overnight and IFNγ positive spots measured **(B).** Proliferation of CD8+ **(C, left)** and CD4+ **(C, right)** T cells was measured by dye dilution after incubation for seven days at 37°C, 5% CO_2_. Error bars represent the mean ± SD. p values were determined by two-tailed, student’s t-test. *\*p < 0.05*.

We next tested NTX250 ex vivo using peripheral blood mononuclear cells (PBMCs) from two patients diagnosed with cervical intraepithelial neoplasia (CIN). A total of 2×10⁵ PBMCs per condition were stimulated with the following mRNA formulations: non-coding control mRNA, LIGHT mRNA, IL-12 mRNA, HPV16 E6-E7 mRNA, or NTX250, each administered at a dose of 2 ng mRNA per 1000 cells. After 24 hours, IFNγ production was assessed by ELISpot. As expected, neither Nutshell™ formulated non- coding mRNA nor LIGHT mRNAs induced detectable HPV16 E6-E7-specific immune responses as measured by IFNγ spot-forming units in ELISpot. In contrast, both Nutshell™ formulated IL-12 and HPV16 E6-E7 mRNAs induced substantial E6-E7-specific IFNγ responses, and NTX250 treatment led to the strongest IFNγ responses in PBMCs from both donors (Figure 5B).

To evaluate the proliferative capacity induced by each formulation, a 7-day proliferation assay was performed in which PBMCs were labeled with a violet tracking dye and treated under the same stimulation conditions. Non-coding control and LIGHT mRNAs failed to induce measurable proliferation, while IL-12 and HPV16 E6-E7 mRNAs triggered only minimal expansion. In contrast and remarkably, NTX250 treatment resulted in robust and consistent proliferation of CD8+ and CD4+ T cells in both CIN patient samples, indicating a superior capacity to activate and expand antigen-specific lymphocytes ex vivo (Figure 5C).

The marked increase in numbers and proliferative activity of functional antigen-specific T cell responses in NTX250-treated samples—beyond what was observed with either IL-12 mRNA or HPV16 E6-E7 mRNA alone— demonstrates a synergistic effect, where antigen specific T-cells are selectively expanded by receiving TCR signaling through the HPV16 E6-E7 antigen, IL12 providing survival and expansion (signal 3) signaling and LIGHT co-stimulation (signal 2).

## Discussion

This study presents NTX250 as a rationally designed, multimodal mRNA-based therapeutic vaccine with potent immunostimulatory activity against HPV16-associated malignancies. Through a combination of subcellularly targeted antigen design and localized immunomodulation, NTX250 addresses longstanding challenges in cancer immunotherapy, namely poor antigen presentation, insufficient T cell priming, and immune suppression within the tumor microenvironment. Our results demonstrate that mNTX250, mouse surrogate of NTX250, not only eradicates large, established HPV16- transformed tumors in murine models but also induces durable immune memory, suggesting potential of NTX250 as a non-surgical therapeutic approach for cervical intraepithelial neoplasia (CIN) and other HPV-driven pathologies.

A key feature of NTX250 is the strategic engineering of the E6-E7 fusion antigen. By targeting the fusion protein to endolysosomal compartments via CD1d trafficking domains, the construct ensures efficient presentation via both MHC class I and II pathways, enhancing CD8+ and CD4+ T cell responses. These findings reinforce the concept that subcellular localization is a critical determinant of antigen processing and immunogenicity, consistent with previous work in tumor and infectious disease vaccine design^20,32,33^. Moreover, utilizing the versatility of mRNA technology, we investigated how altering the subcellular localization of mRNA-encoded proteins, such as LIGHT, can be employed to enhance their functional activity. This approach facilitated the design of a membrane-stabilized engineered LIGHT, which demonstrated improved HVEM-mediated signaling and immune activation, as detailed in the supplementary information and Supplementary figure 6.

Our study also highlights the synergistic function of IL-12 and LIGHT as immunomodulators. IL-12 promotes Th1 differentiation and IFN-γ secretion^34–36^ while LIGHT enhances co-stimulation, T cell recruitment and myeloid activation through HVEM and LTβR pathways^37–41^.

The therapeutic efficacy of NTX250 was demonstrated even at low doses, as evidenced by complete tumor regression and memory response induction across multiple dosing levels. Importantly, NTX250 outperformed formulations lacking either IL-12 or LIGHT, and its effectiveness was abrogated by CD8+ T cell depletion, affirming the central role of cytotoxic T lymphocytes in mediating tumor clearance. The observed reshaping of the TME—with increased infiltration of CD8+ T cells, NK cells, and dendritic cells, alongside a marked reduction in regulatory T cells and a Th1-skewed cytokine milieu— provides a mechanistic explanation for the vaccine’s potent antitumor effects. In contrast to existing HPV-targeted therapeutic vaccines such as VGX-3100 ^42^, which rely on intramuscular DNA delivery and elicit modest regression of high-grade lesions, NTX250 demonstrated complete tumor eradication in preclinical models, with evidence of robust and durable memory responses. While systemic delivery of mRNA vaccines, as exemplified by the lipoplex formulation of Grunwitz et al., can elicit anti-tumor immunity, NTX250 is designed for intra-lesional administration, enabling a higher local concentration of both antigen and immunomodulatory signals; this targeted delivery, combined with the co-expression of IL-12 and LIGHT, contributes to the observed complete tumor eradication and long-lasting immunity in preclinical models^43^.

Moreover, NTX250 demonstrated translational potential through its ability to elicit HPV16- specific immune responses in non-human primates and PBMCs derived from CIN patients. These responses were stronger than those induced by control formulations or individual immunomodulators, suggesting that the synergistic integration of antigen, IL-12, and LIGHT is required for maximal immunogenicity. The ex vivo human data supports the feasibility of using NTX250 in a therapeutic setting for HPV-infected individuals at risk of progression to cervical cancer. Compared to previously described mRNA vaccines that encode tumor antigens alone^44^, the multi-component design of NTX250 offers a more comprehensive approach to immune activation and tumor microenvironment remodeling. Additionally, in a model involving established subcutaneous MC38 tumors, animals treated with mNTX255 (mRNA formulation designed on NTX250 platform encoding mIL-12, mLIGHT, and an antigen of MC38 neoepitopes) demonstrated significant tumor growth inhibition, with tumor control being similar between animals treated with mNTX255 and mNTX151. Spleen analyses showed a marked increase in T cells specific for multiple neoepitopes in the mNTX255 group, confirming that the vaccine effectively elicited targeted immune responses in vivo. These findings highlight the capacity of NTX250 platform to generate neoepitope-specific T cell immunity, a critical feature for personalized cancer immunotherapy.

While IL-12 is a well-characterized immune stimulant with demonstrated anti-tumor efficacy, its clinical development has been severely hampered by systemic toxicity. Early-phase trials of recombinant IL-12 revealed significant adverse events, including severe cytokine release syndrome and treatment-related mortality reported in a phase II trial for renal cell carcinoma, which led to a temporary clinical hold^45,46^. These toxicities were largely attributed to widespread systemic exposure and uncontrolled IFN- γ induction. In contrast, NTX250 is formulated to achieve localized cytokine expression, and pharmacokinetic profiling in preclinical models revealed that IL-12 encoded by NTX250 is present at concentrations over 100-fold higher in the injected tumor relative to circulation. This pronounced localization likely mitigates the risk of systemic IL-12–related toxicity^22^ while preserving its potent immunostimulatory function within the tumor microenvironment. This compartmentalized cytokine delivery strategy may represent a significant advance in the safe clinical application of IL-12–based immunotherapies.

Taken together, our findings suggest that NTX250 is a promising therapeutic candidate capable of orchestrating robust, localized, and durable anti-HPV immunity through synergistic molecular components and rational design. Future studies will focus on the clinical translation of NTX250, including dose optimization, safety profiling, and assessment of efficacy in patients with high-grade CIN and other HPV-related malignancies. NTX250 represents a new class of precision immunotherapeutics that integrates advanced mRNA engineering, immune co-stimulation, and TME remodeling to overcome critical barriers in the treatment of virally driven cancers.

## Acknowledgements

This study was performed with help from the Beckman Center for Immune Monitoring/FCIM core at USC that is supported by a NIH grant P30 CA014089. W. Martin Kast holds the Walter A. Richter Cancer Research Chair

## IRB and IACUC approvals

Patients were required to comprehend and sign an informed consent form approved by the Institutional Review Board (IRB) of LAC+USC Medical Center (IRB# HS-10-00489). The animal protocol and all procedures at USC were approved by the USC Institutional Animal Care and Use Committee (Permit number 20065).

## Author Contributions

Conceived and designed the experiments: OH, SD, ML, DMD, WMK. Performed the experiments: OH, ML, WL, AS, NF, OL, CR, AS, CM, SN, NP, EW, XW, PD, EL, AS, DMD, RP and DJF. Analyzed data: OH, SD, ML, NF, OL, WL, CR, DMD, WMK. Wrote the initial draft of the manuscript: OH, SD, ML. All authors critically read, edited and approved the final manuscript.

## Material and Methods

### mRNA synthesis and Nanoparticle formulations

The DNA templates were amplified from plasmids with Platinum SuperFi II DNA polymerase (Invitrogen). DNA templates were purified using RNAClean XP (Beckman Coulter) magnetic beads. The mRNA was synthesized from DNA template by in vitro transcription using HiScribe T7 RNA polymerase (NEB), ATP/CTP/GTP (NEB), N1-methylpseudouridine-5ʹ-triphosphate (TriLink). The mRNA was co-transcriptionally capped using 9.5 mM CleanCap AG (TriLink). The reaction was incubated at 37◦C for 4h followed by treatment with TURBO DNase (Invitrogen). Post-DNase treatment, the mRNA was purified using cellulose chromatography (Cellulose Type 50, Sigma Aldrich) and RNAClean XP magnetic beads. mRNA yield was quantified using Nanodrop One, and purity was determined using Fragment Analyzer (Agilent Technologies; RRID: SCR_019411). mRNA cargos were formulated for in in vivo and in vitro experiments using Nutshell^TM^ DVs as previously described^11^. Briefly, mRNAs in pH 5 citrate buffer were mixed on a microfluidic device with Nutshell^TM^ lipids, DSPC, cholesterol, and DMG-PEG2000 in ethanol. The resulting particles were dialyzed against a citrate-sucrose buffer and stored at − 80◦C until use.

### CIN antigen specific responses and proliferation in human PBMCs

PBMCs were obtained from Charles River Labs, STEMCELL technologies and AllCells. CIN patient PBMCs were collected in the lab of Prof. Martin Kast at USC under IRB protocol # HS-10-00489^47^. For CIN patients, leukapheresis was performed on blood samples to enrich peripheral blood mononuclear cells (PBMC), which were additionally purified immediately after collection using Lymphocyte Separation Media (Corning) by density gradient centrifugation, and cryopreserved in liquid nitrogen.

For T cell proliferation, 1 × 10^4^ cells/well HEK293 cells were transfected in 96-well flat-bottom plates with mRNA encoding hIL-12 and hLIGHT either alone or in combination in Lipofectamine Messenger Max according to the manufacturer’s protocol. Control cells were transfected with noncoding control mRNA. CD3+ T cells were then isolated from PBMCs from CIN donors using the EasySep^TM^ Human T Cell Isolation Kit according to the manufacturer’s instructions (Miltenyi). Purified CD3+ T cells were resuspended at 5 × 10^6^ cells/mL and labeled with CellTrace^™^ Violet dye according to the manufacturer’s protocol.

Half of CD3+ T cells were then treated with anti-human CD3 antibody (200 ng/mL), the other half was left untreated. 100 μL of the resulting cell suspensions were then added to HEK293 cell cultures (1 × 10^5^ cells/well) generating + or – anti-CD3 treated T cells conditions for every mRNA treated HEK293 condition and incubated for 4 days at 37°C and 5% CO_2_. The final concentration of anti-CD3 antibody in cell cultures was 100 ng/mL. After 4 days, cells were harvested, and T cell proliferation was measured by flow cytometry on Northern Lights cytometer (Cytek). Fold increase in proliferation is measured relative to untreated CellTrace^TM^ Violet dye-stained cells for which the dye intensity stays constant.

For human IFNγ ELISpot, donor PBMCs were thawed and seeded at 1.5 × 10^6^ cells/mL in a 24-well tissue culture-treated plate and incubated overnight at 37°C, 5% CO_2_. The next day, cells were treated with either HPV16E6E7-hCD1d mRNA or noncoding mRNA control in Nutshell^TM^ formulations. At 4 hours post-treatment, 3 × 10^5^ cells per condition were transferred to an enzyme-linked immunosorbent spot (ELISpot) plate precoated with anti-IFN-γ capture antibody (BD Bioscience) and incubated at 37°C, 5% CO_2_ for 24 hours. The remaining 1.2 × 10^6^ cells were left in the original 24-well plate and incubated at 37°C, 5% CO_2_ for 18 hours. The ELISpot plate was developed to assess formation of IFN-γ-positive spots using the Immunospot plate reader and software (CTL Ltd., Cleveland, OH).

### Mice

For in vivo studies with C3.43 tumors, pathogen-free 5-8 week old C57BL/6 female mice were purchased from Taconic and housed at University of Southern California (USC). Studies were performed in compliance with and approved by USC Institutional Animal Care and Use Committee (IACUC). For in vivo studies with MC38 tumors, 6-8 week old C57BL/6 mice were purchased from Charles River Laboratories and housed at Charles River Accelerator and Development Lab (CRADL), and studies were performed in compliance with CRADL IACUC. Blood was collected in-life by retro-orbital or cheek bleeds or following euthanization by cardiac puncture into K2-EDTA microtainer. Cells were centrifuged at 300 x g for 5 minutes at room temperature and plasma collected. Cell pellets were incubated with red blood cell lysis buffer and washed with PBS to prepare PBMCs for further analysis.

### In vitro cell culture

C3.43, a progressive subclone of the HPV16-transformed B6 mouse embryo cell line C3, were used for tumor challenge studies. C3.43 cells were grown and expanded in vitro with IMDM medium supplemented with heat-inactivated 10% fetal bovine serum (Gemini), 50 µM 2-ME, and 50 µg/mL gentamicin. TC-1 cells were cultured and expanded in vitro with RPMI-1640 medium supplemented with 10% fetal bovine serum (FBS; Omega Scientific, Riverside, MO), 2 mM Lglutamine, 50 µg/ml kanamycin, 50 mM 2-mercaptoethanol, 1 mM sodium pyruvate, 2 mM nonessential amino acids, 0.4 mg/ml G418, and 0.2 mg/ml hygromycin. HeLa, MC38, and THP-1 were cultured in RPMI 1640 medium containing 10% fetal bovine serum (FBS, Gibco), 100 international units/mL penicillin, 100 μg/mL streptomycin (Invitrogen). Cells were cultured at 37 °C in 5% CO_2_.

To generate human monocyte-derived dendritic cells (mo-DCs), monocytes were isolated from human PBMCs (StemCell technologies) using EasySep Human Monocyte Isolate kit (StemCell) according to the manufacturer’s instructions. Monocytes were plated in non-tissue culture treated sterile petri dishes in RPMI media (Gibco) containing 10% FBS, 1x Antibiotic-Antimycotic (Gibco) and 50ng/mL GM-CSF and 25 ng/mL IL-4 at 37 °C in 5% CO_2_. On day 3, complete media including GM-CSF and IL-4 was replaced. On day 6, cells were harvested using ice-cold PBS and counted before use.

### Tumor challenge

Mice were challenged subcutaneously with 1 × 10^5^ C3.43 cells or 1 × 10^6^ MC38 cells. Prior to first vaccination, tumors were measured with manual calipers by measuring tumor length, height, and depth to generate a tumor volume (Vol = l × w × d).and mice were randomized into study groups. For TC-1 tumor cell re-challenge study, 21 weeks after initial tumor implantation 26 surviving mice and 10 B6 naïve control mice were challenged subcutaneously on left flank with 1 × 10^5^ TC-1 tumor cells in 100 µL HBSS. For CD8 T cell depletion, mice were administered anti-mCD8 antibody (clone 2.43; BioXCell) or isotype control (BioXCell) intraperitoneally for 3 consecutive days starting 89 days post initial tumor challenge. Once CD8 depletion was confirmed, mice were rechallenged with 1×10^5^ C3.43 cells subcutaneously on the opposite flank. Depletion was maintained with twice weekly administration of anti-CD8 or isotype control for duration of tumor rechallenge. Mice were monitored daily, and tumor volume was measured twice weekly. Mice were sacrificed if the tumor reached a maximum allowed volume of approximately 1,500 mm^3^ or showed a maximum permissible level of discomfort.

### Flow Cytometry

The following fluorochrome-conjugated rat anti-mouse mAbs (Biolegend) were used for flow cytometry: CD3e-FITC, CD8-PerCP-Cy5.5, CD4-APC, CD11b-PE-Cy7, CD19-PE-Cy7, NK1.1-BV411, FOXP3-PE. Tetramer staining was done with PE-labeled H-2Db HPV16 (RAHYNIVTF). The following anti-human mAbs were used: by CD3-PerCP-Cy5.5, CD4-BV650, CD8-FITC and CD25-PE. Antibodies were purchased from BD Biosciences, Invitrogen, Biolegend or eBioscience. Cells were stained with Zombie Aqua^TM^ or Zombie NIR^TM^ for viability, then surface marker staining was done for 20 min at room temperature in PBS (Gibco) containing 0.01% sodium azide (Sigma Aldrich) and 0.5% BSA (Sigma). Intracellular staining for FoxP3 was performed according to manufacturer protocol (BioLegend).

For expression of E7, HeLa cells were transfected with mRNA encoding different antigen designs using Lipofectamine MessengerMax transfection reagent according to the manufacturer’s instructions. For surface expression, mRNA transfected cells were stained with mouse anti-E7 antibody (Clone 8C1 Thermo Scientific) 24 hours after transfection for 20 minutes at 4°C, followed by incubation with anti-mouse IgG-PE secondary antibody. For intracellular expression of E7 protein, cells were fixed and permeabilized using BD perm/wash buffer (BD Biosciences) for 30 minutes at 4°C prior to incubation with mouse anti-E7 antibody and anti-mouse IgG-PE secondary.

Stained cells either were fixed in 2% paraformaldehyde or were run fresh and were analyzed by flow cytometry on a FACSCanto II (BD Biosciences) or Northern Lights^TM^ (Cytek Biosciences). Data were analyzed using FlowJo software.

### Preparation of mouse peripheral mononuclear cells (PBMCs) and plasma

Blood was collected in-life by retro-orbital or cheek bleeds or following euthanization by cardiac puncture into K2-EDTA microtainer. Cells were centrifuged at 300 × g for 5 minutes at room temperature and plasma collected. Cell pellets were incubated with red blood cell lysis buffer and washed with PBS to prepare PBMCs for further analysis.

### IFNγ ELISpot

For human IFNγ ELISpot, donor PBMCs were thawed and seeded at 1.5 × 10^6^ cells/mL in a 24-well tissue culture-treated plate and incubated overnight at 37°C, 5% CO_2_. The next day, cells were treated with either HPV16E6E7-hCD1d mRNA or noncoding mRNA control in Nutshell^TM^ formulations. At 4 hours post-treatment, 3 × 10^5^ cells per condition were transferred to an enzyme-linked immunosorbent spot (ELISpot) plate precoated with anti-IFN-γ capture antibody (BD Bioscience) and incubated at 37°C, 5% CO_2_ for 24 hours. The remaining 1.2 × 10^6^ cells were left in the original 24-well plate and incubated at 37°C, 5% CO_2_ for 18 hours. The ELISpot plate was developed to assess formation of IFN-γ-positive spots using the Immunospot plate reader and software (CTL Ltd., Cleveland, OH). For mouse ELISpot, spleens were processed to single cell suspension and plated at 2 × 10^5^ per well and stimulated overnight with 1 µM peptides for Adpgk (ASMTNMELM), Cpne1 (SSPYSLHYL), Reps1 (AQLANDVVL) or Rpl18 (KAGGKILTFDRLALESPK), HPV16 E7_49-57_ (RAHYNIVTF), PADRE (AKFVAAWTLKAAA) (ELIM Bio), or overlapping peptides for HPV16 E6 and E7 (JPT Peptide Technologies GmBH). Assay performed using the BD Biosciences or MabTech mouse IFNγ kit according to the manufacturer’s instructions, and IFNγ spot-forming cells were measured using an ImmunoSpot (CTL) or ASTOR^TM^2 (MabTech) analyzer and software.

### Isolation of tumor supernatant and Tumor-Infiltrating Lymphocytes (TILs)

Tumors were dissociated using mouse tumor dissociating enzymes (Miltenyi) using GentleMACS (Miltenyi) with incubation at 37°C for 40 minutes. Dissociated tumor samples were centrifuged to separate tumor supernatant and tumor cells. Tumor cells were filtered, and samples were frozen before analysis.

### Cytokine analysis

Cytokines in plasma and tumor supernatant samples were assessed using mouse ProCarta 48-plex kit (Luminex) according to the manufacturer’s instructions. Samples were run on MagPix and analyzed with xPONENT software (Luminex).

### Immune responses in nonhuman primates (NHPs)

Cynomolgus macaques (male) were dosed intramuscularly with 375 µg (n=2) or 1125 µg (n=2) of NTX250 on days 0, 7, and 14. EDTA-blood was collected on days -7, 7, and 21 and processed to PBMCs using Leucosep^TM^ tubes (MP Bio). This study was conducted under protocols approved by the institutional animal care and use committee overseeing animal welfare at Biomere/JOINN laboratories.

2 × 10^5^ PBMCs were incubated anti-human IFNγ-coated ELISpot plates (BD Bioscience) with an overlapping peptide libraries for HPV16 E6 and E7 (JPT Peptide Technologies GmBH) in overnight at 37°C with 5% CO_2_ and developed using alkaline phosphatase detection reagents (Invitrogen). Cells were incubated without peptide as a negative control. Plates were scanned and quantified using the Immunospot plate reader and software (CTL Ltd., Cleveland, OH).

### TCR reporter assay

HEK293 cells were seeded at a density of 40,000 cells per well in 100 µL of Opti-MEM media in a round-bottom 96-well cell culture plate. Following cell attachment, the cells were treated with increasing amounts of mRNA formulated in Messenger MAX™ (Thermo Fisher Scientific), ranging from 0.675 ng per 1,000 cells with a 2-fold increase up to 5 ng per 1,000 cells, delivered in 50 µL of Opti-MEM/Messenger MAX™. The cells were incubated at 37°C in a humidified atmosphere containing 5% CO2 for 24 hours. After the 24-hour incubation, the media was replaced with DMEM supplemented with 10% fetal bovine serum (FBS) and 1% penicillin-streptomycin. Transfected cells were incubated at 37°C + 5% CO2 overnight. The following day, Jurkat-Lucia NFAT reporter lines stably expressing CD8 and a TCR with specificity for HLA-A:02 epitopes E7_11-20_ (BPS Bioscience, clone 1G4, Ca#78675) cultured in RMPI (Gibco, Ca# 11875-093)), 10% FBS (Gibco, Ca# A5209502), 2-Meraptoethanol (Gibco Ca# 21985-023) and Penicillin-Streptomycin (Gibco, Ca# 15-140-122) were added to Jurkat-Lucia NFAT reporter line cell suspension. Jurkat-Lucia NFAT reporter cells and transfected HEK293 cells were co-cultured for approximately 24hrs at 37°C + 5% CO_2_. 20ul cell culture supernatant was collected and mixed with 50ul QUANTI-Luc™ 4 Lucia/Gaussia luciferase detection reagent (Invivogen, Ca# rep-qlc4lg5). Luminescence was detected on a SpectraMax iD3 spectrophotometer (Molecular Devices).

### LIGHT expression and activity

For soluble and membrane expression of LIGHT, HEK293 cells were transfected with mRNA encoding wildtype or engineered LIGHT using Lipofectamine MessengerMax transfection reagent (Invivogen) and after 24 hours, supernatants and cells were harvested. For membrane expression, transfected cells were incubated with recombinant human LTβR protein (R&D Sytems) for 30 minutes at room temperature, followed by incubation with anti-human IgG-Fc-PE antibody (Biolegend) for 30 minutes at room temperature. Soluble LIGHT was quantified by ELISA using TNFSF14 ELISA kit (MyBiosource). LIGHT activity was measured using an HVEM reporter cell line consisting of Jurkat cells stably expressing HVEM and signal-dependent luciferase gene (BPS Bioscience). HVEM reporters were incubated with supernatants or cells following mRNA transfection and luciferase signal measured as relative light units (RLU) after 6 hours.

### LIGHT co-stimulation assay

A flat bottom 96 well TC-treated plate was coated with anti-human CD3 (aCD3; BioLegend clone OKT3 Ca# 317326) in 100µl volume at 1µg/ml in PBS. The coated plate was stored overnight at 4°C, then washed and coated with recombinant human LIGHT (BioLegend Ca# 762402) in 100µl volume with a two-fold dose titration in PBS with starting concentration at 20 µg/ml. The plate was incubated at 37°C for 4hrs.

T cells were isolated from donor PBMCs using a STEMCELL human T cell isolation kit (STEMCELL Ca# 17951). T cells were resuspended in PBS at a concentration of 5 × 10^6^ cells/mL and labeled with CellTrace^TM^ Violet (Thermo Ca# C34557) at a 1:1000 volumetric ratio for 20 minutes at 37°C + 5% CO_2_. The cell suspension was quenched with media, washed and resuspended in complete AIM-V/RPMI media to a concentration of 1 × 10^6^ cells/mL.

The 96 well tissue culture plate with immobilized anti-CD3 and recombinant LIGHT was washed with PBS and 1 × 10^5^ CellTrace^TM^ Violet-labeled T cells were added per well in 100µl volume. Soluble recombinant human LIGHT (BioLegend Ca# 762402) or anti-human CD28 (aCD28; BioLegend clone CD28.2 Ca# 302934) was added to either 20µg/mL or 5µg/mL final volume and final volume was adjusted to 200µl/well for all samples. Cells were incubated at 37°C + 5% CO_2_.

The following day, T cells were evaluated for proliferation by CellTrace^TM^ Violet signal diffusion and activation by CD25 upregulation. Cells were stained with Zombie NIR^TM^ for viability, followed by PerCP Cy5.5 anti-human CD3, BV650 anti-human CD4, FITC anti-human CD8 and PE anti-human CD25. Flow cytometry was performed on a Cytek Northern Lights^TM^ system and data was evaluated with FlowJo^TM^ software.

### Confocal Microscopy

Cell line: HeLa cells (CCL-2, ATCC, Manassas, VA, USA) derived from cervical carcinoma were maintained in Iscove’s Modified Dulbecco’s Medium (Gibco, Waltham, MA, USA) supplemented with fetal bovine serum (10%; Omega Scientific, Tarzana, CA, USA), 2-mercaptoethanol (0.05 mM; Gibco), and gentamycin (50 units/mL; Gibco) at 37 °C with 5% CO2 and 95% relative humidity. Antibodies: Anti-human papillomavirus type 16 E7 (28-0006), anti-SERCA2 ATPase (JM10-20), and anti-VPS25 (PA5-99005) were purchased from ThermoFisher Scientific (Waltham, MA, USA). The secondary antibodies Texas Red-X goat anti-rabbit (T6391) and Alexa Fluor 488 goat anti-mouse (A11029) were also purchased from ThermoFisher Scientific. Anti-EEA1 (ab2900) and Golgin97 (ab84340) were purchased from Abcam (Cambridge, UK). LAMP1 (D2D11) was purchased from Cell Signaling Technology (Danvers, MA, USA). Immunofluorescence microscopy: For HPV16 E7 colocalization with organelle markers, cells were seeded on 8-well chamber slides with #1.5 polymer coverslip bottoms and ibiTreat surface modification for improved cell attachment (Ibidi, Fitchburg, WI, USA) and grown overnight. Cells were incubated with E6- E7 constructs for 1, 2, 4, or 6 hours at 37 °C prior to fixation with 4% paraformaldehyde for 10 minutes at room temperature. Cells were permeabilized in DPBS containing 0.1% Triton X-100 for 10 minutes and subsequently blocked for 1 hour in DPBS containing 1% bovine serum albumen. Cells were stained overnight for HPV16 E7 (1:100) and either EEA1 (5µg/ml), SERCA2 (1:100), VPS25 (1:100), Golgin97 (5µg/ml), or LAMP1 (1:100). Cells were secondarily stained with antibodies labelled with either Alexa Fluor 488 or Texas Red-X and counterstained with Hoescht 33342. Images were visualized on a Nikon Eclipse Ti-E Laser scanning confocal microscope. For colocalization analysis, Z-stack images were analyzed using Imaris (Oxford Instruments). E6-E7 construct preparation: Constructs were shipped at 2-8 °C to IMC Lab prior experimentation. Upon receipt, the constructs were visually assessed for quality and subsequently stored at 2-8 °C. On the day of use, constructs were gently flicked to resuspend nanoparticles. Constructs were prepared at a concentration of 0.1ng E6-E7 construct/10k cells, which was experimentally determined to provide optimal signal intensity in immunofluorescence imaging.

