## Supplemental Information for "NTX250: A Modular mRNA-Based Immunotherapy Platform for HPV-Associated Cancers with Broad Applicability"

Supplementary Figure 1

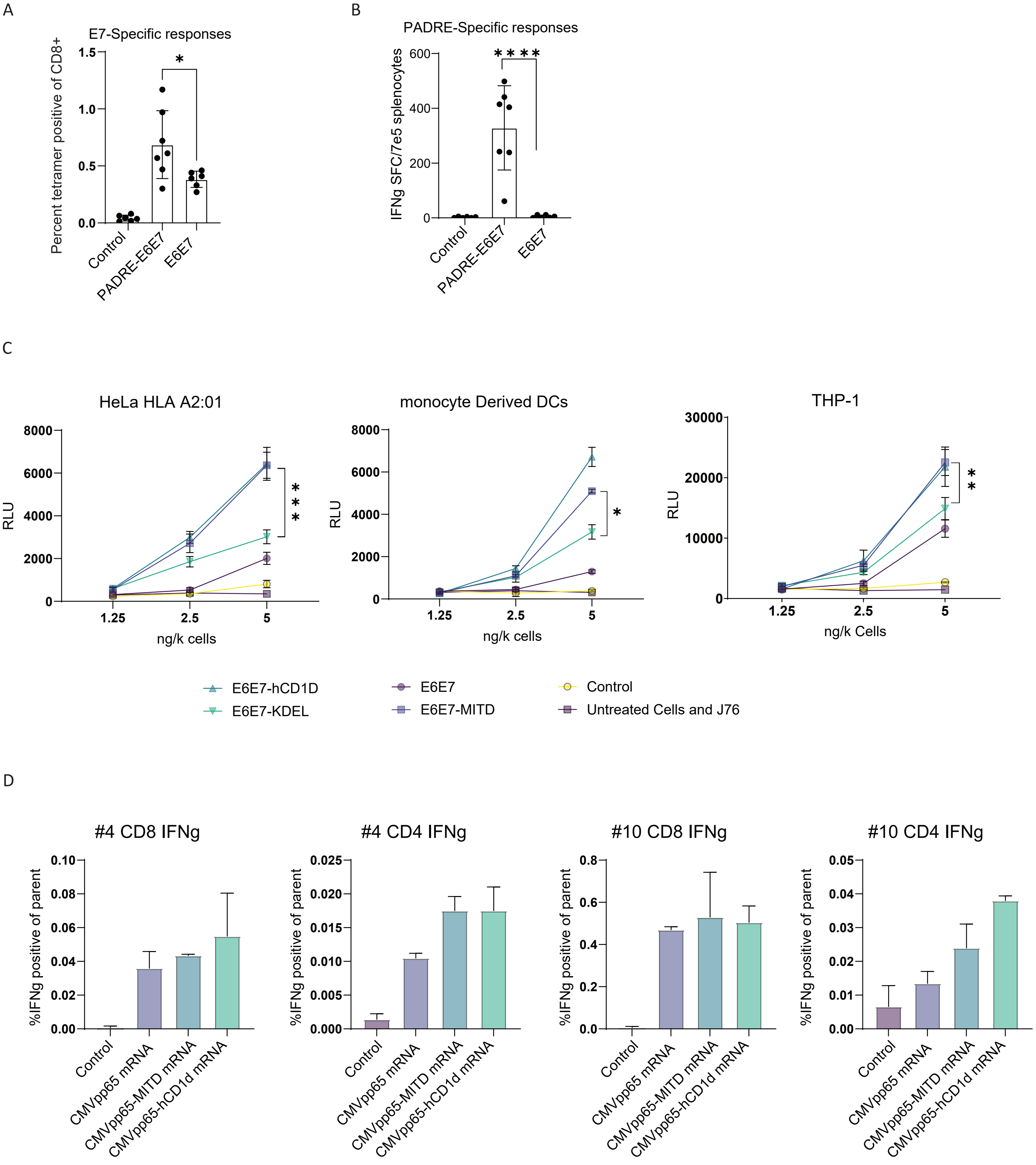

Supplementary Figure 1

E

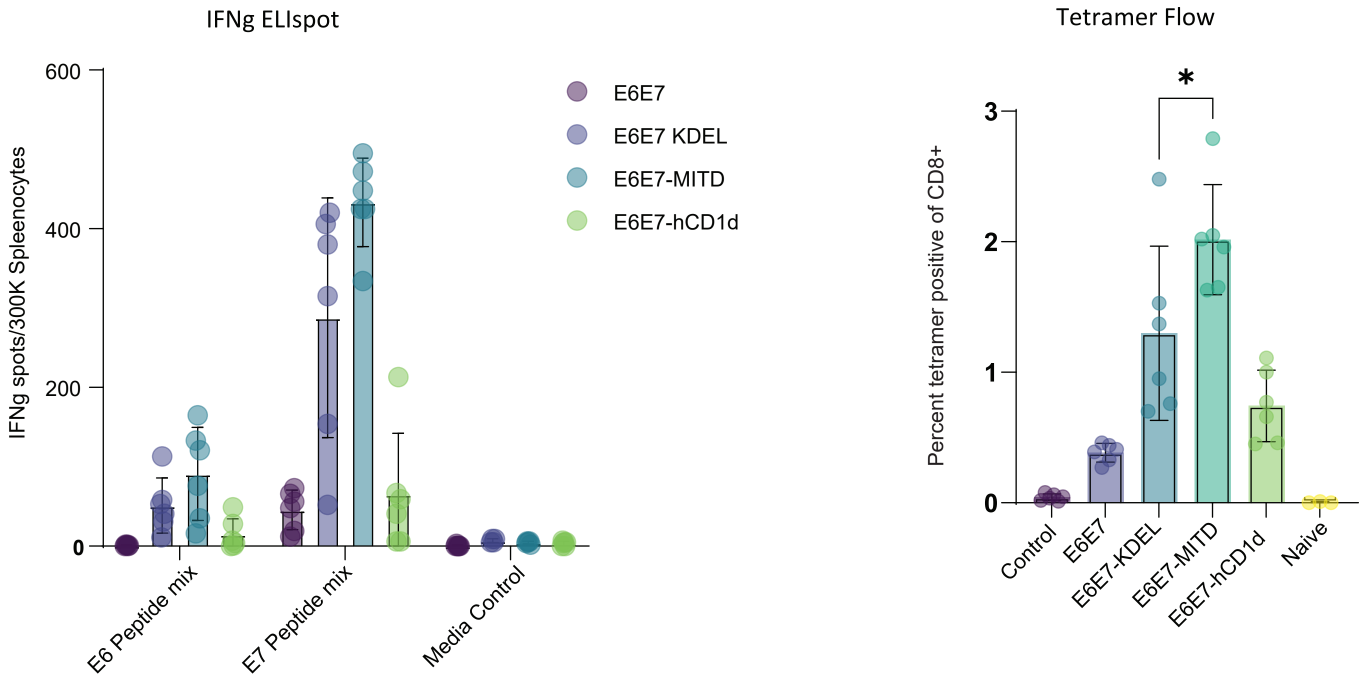


F

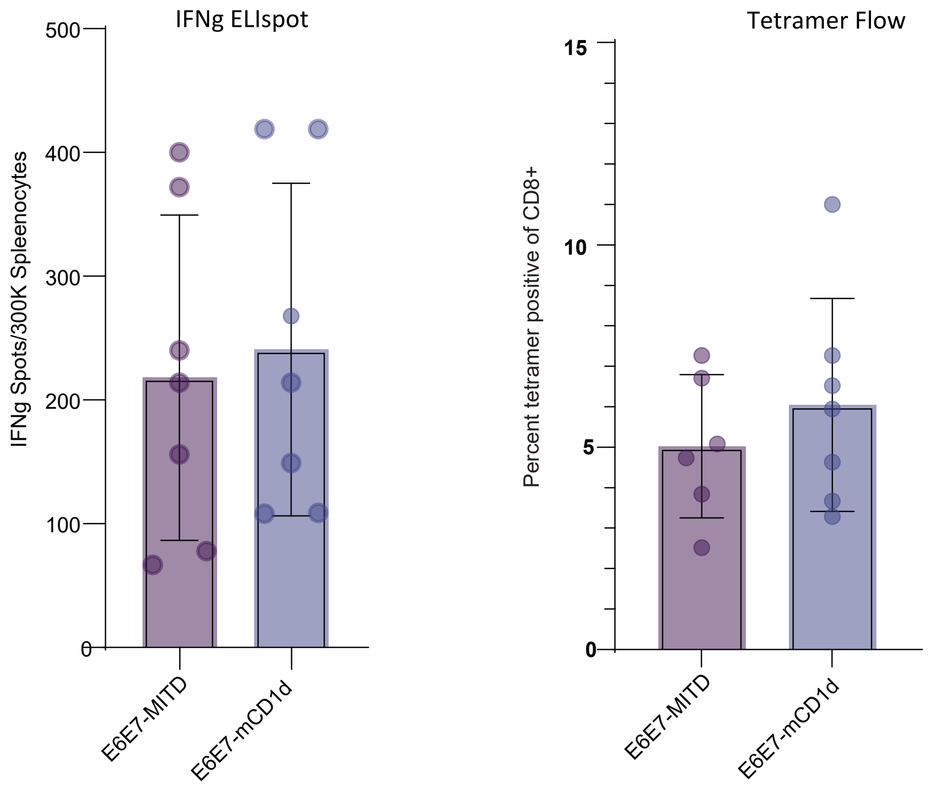

Supplementary Figure 2

A

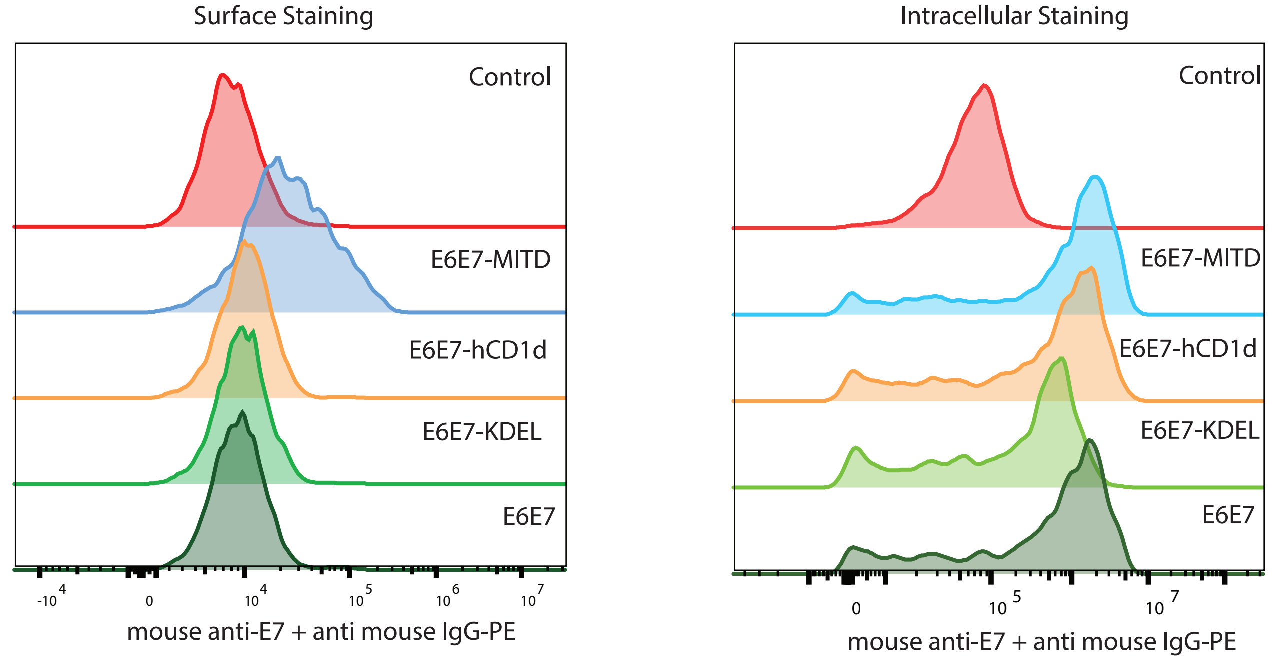


B

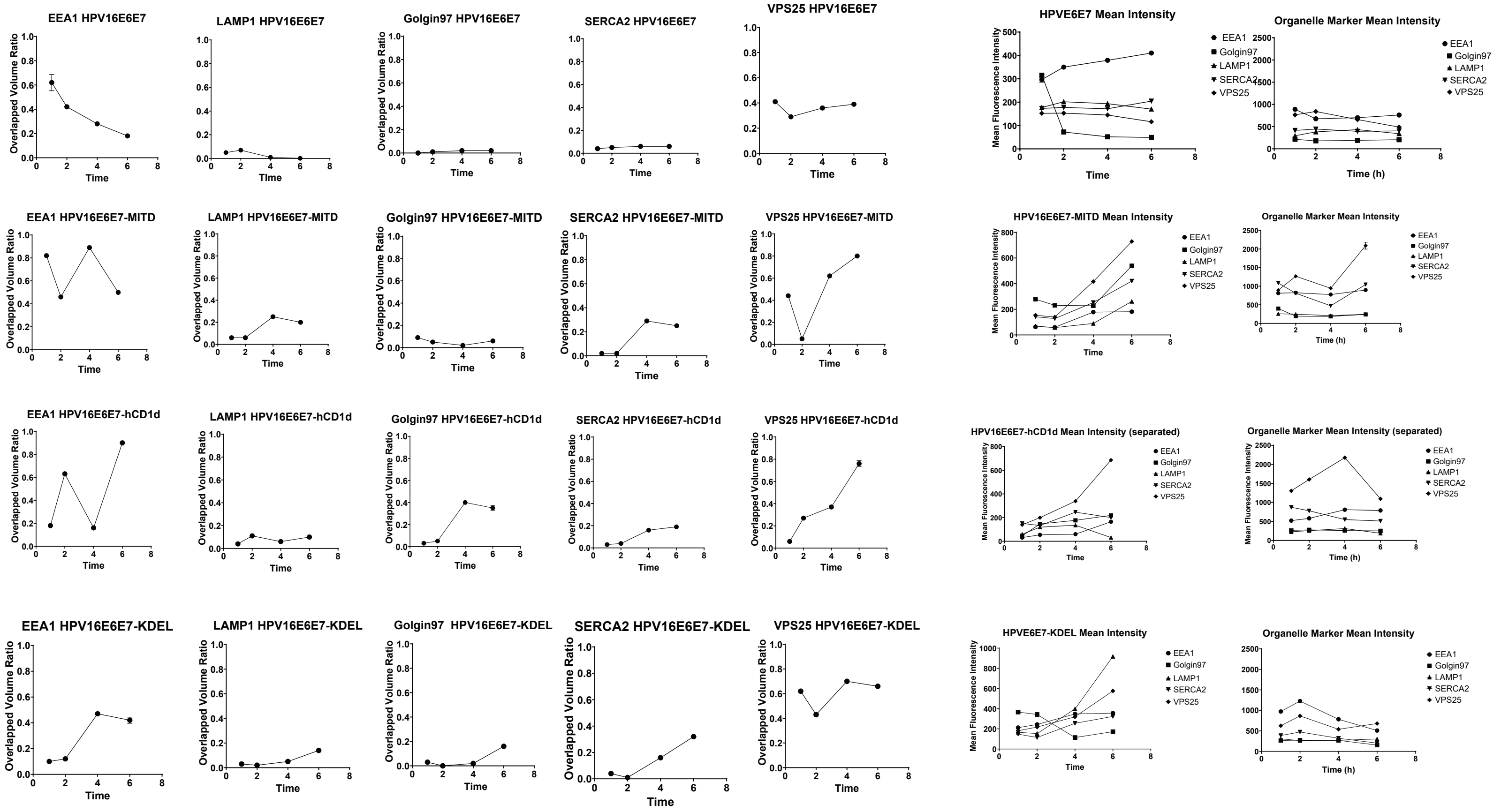

Supplementary figure 3

A

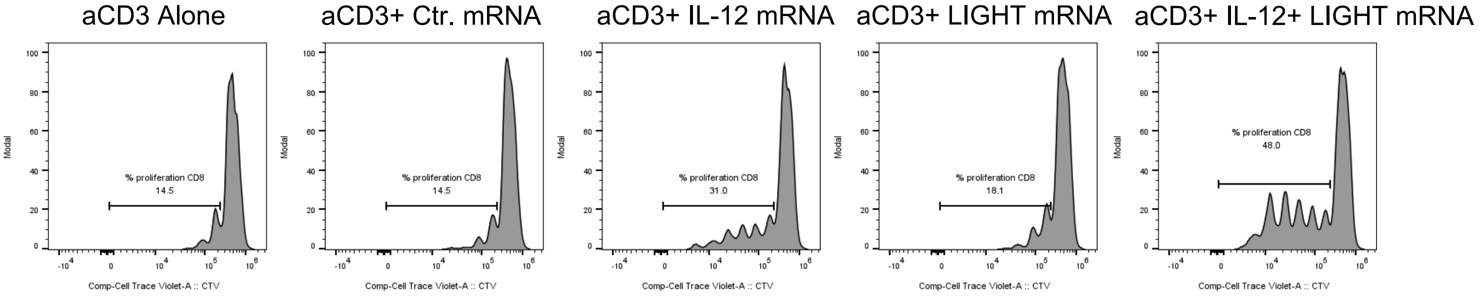


B

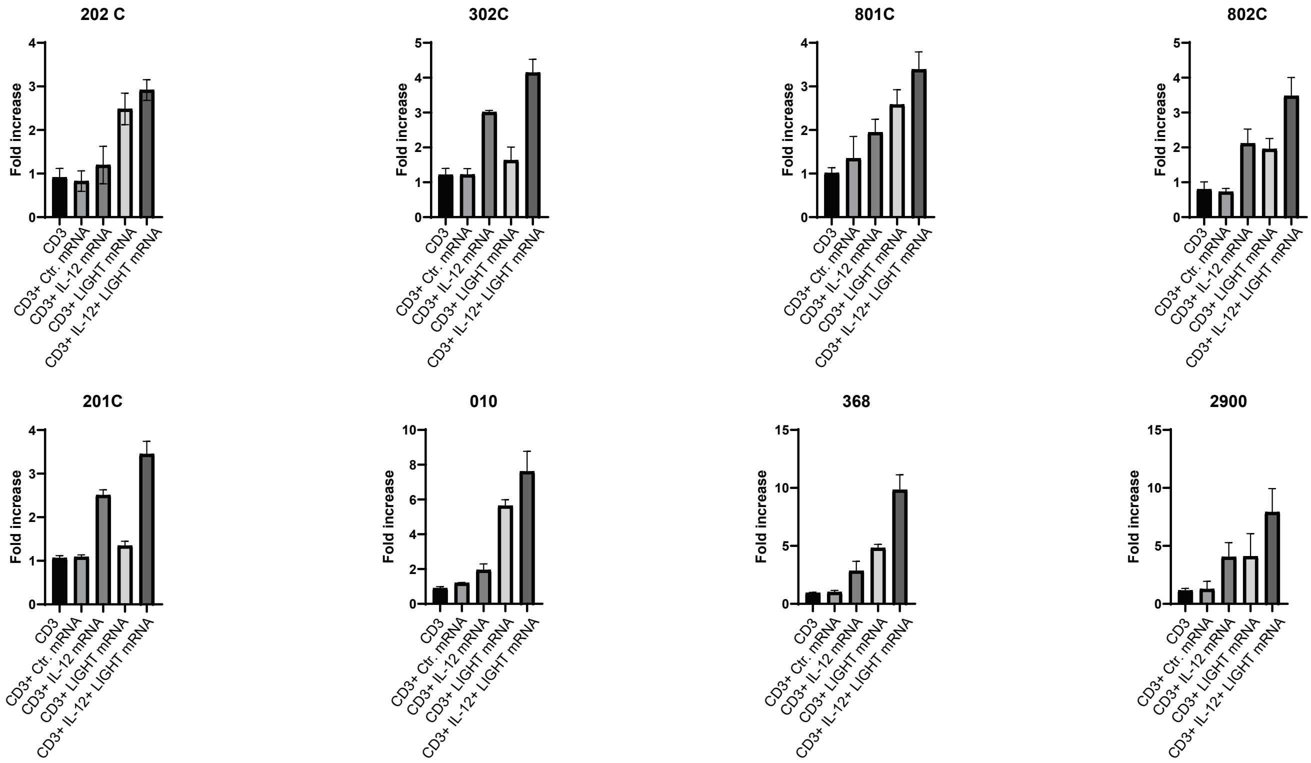

Supplementary Figure 4

A

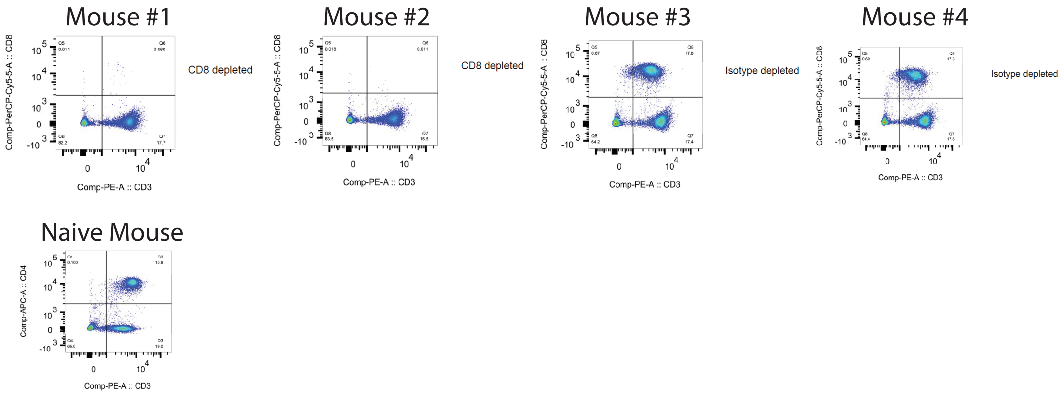


B

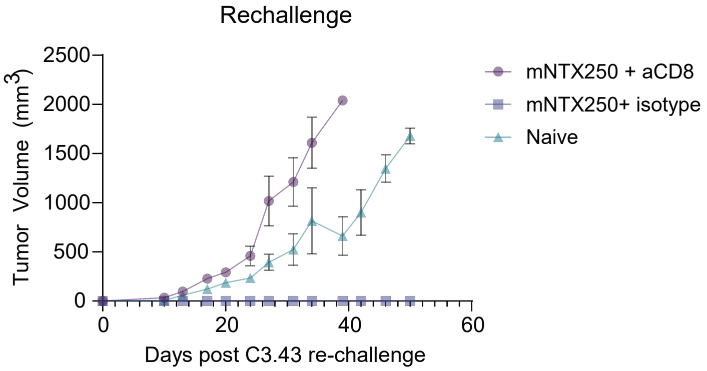

Supplementary Figure 5

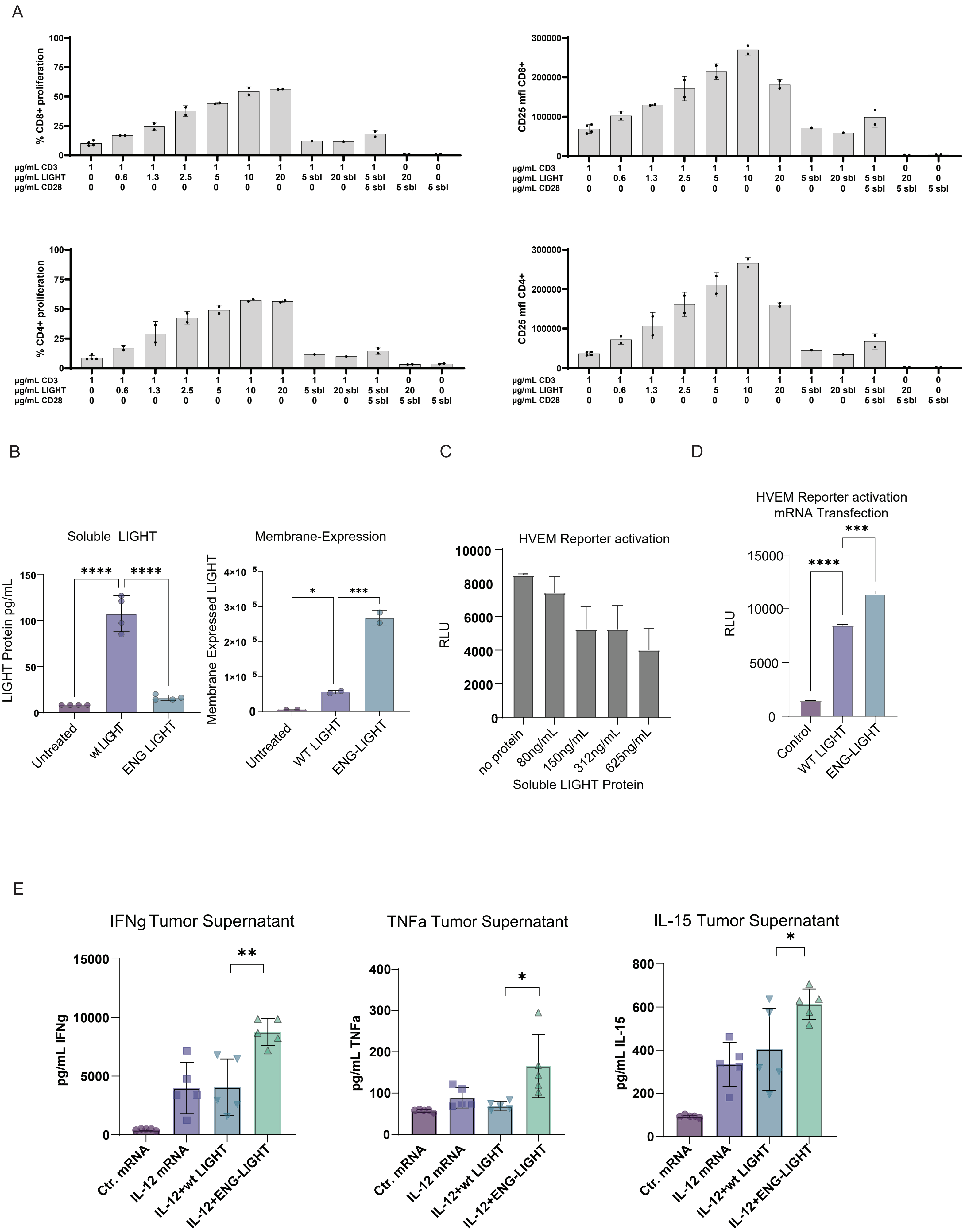

Supplemental figure legends:

**Figure S1: Optimization of HPV16 E6-E7 antigen design. A, B, E, F.** C57BL/6 mice (n=6-7) were immunized twice intramuscularly with 2µg of Nutshell™ formulations of mRNA encoding HPV16 E6-E7 designs with PADRE (PADRE-E6E7) or without (E6E7) or non-coding mRNA (NC mRNA). Seven days after second dose, immune responses were measured by $E7_{49-57}$ tetramer from PBMCs **(A)** or by ELISpot as IFNγ spot-forming units (SFU) for splenic responses against PADRE **(B)**. **C.** HLA-A2+ HeLa, THP-1, and Mo-DCs were transfected with HPV16 E6-E7 with no scaffold (E6E7), hCD1d (E6E7-hCD1d), KDEL (E6E7-KDEL), MITD (E6E7-MITD) or control mRNA at 1.25, 2.5, or 5ng/1000 cells and co-cultured with HPV16 $E7_{11-21}$-specific TCR Tg Jurkat reporter cells. Presentation of HLA-A2-restricted epitope $E7_{11-21}$ was measured by lucia reporter activity (RLU) after 48 hours post transfection (hpt). **D.** PBMCs from two CMV+ human donors (Donor #4 and #10) were transfected with Nutshell™ formulations of mRNA encoding CMVpp65 epitope with no trafficking domain (CMVpp65), MITD (CMVpp65-MITD), or hCD1d (CMVpp65-hCD1d) or control (Ctr) mRNA. IFNγ production from CD4+ and CD8+ T cells were assessed 12 hours post transfection. **E, F.** C57BL/6 mice (n=6-7) were immunized intramuscularly two times separated by two weeks with 2µg of Nutshell™ formulations of mRNA encoding HPV16 E6-E7 with no scaffold domain (E6E7), KDEL (E6E7-KDEL), MITD (E6E7-MITD), hCD1d (E6E7-hCD1d), mCD1d (E6E7-mCD1d) or non-coding mRNA (NC mRNA). Seven days after second dose, immune responses were measured by IFNγ ELISpot as spot-forming units for splenic responses using overlapping peptide pools for HPV16 E6 or E7 or media control **(E, left panel)** or HPV16 $E7_{49-57}$ peptide **(F, left panel)** or by $E7_{49-57}$ tetramer from PBMCs **(E, F, right panels)**. p values were determined by two-tailed, student's t-test. **p < 0.05.**P<0.01. ***p>0.001. ****p>0.0001*

**Figure S2: Subcellular localization of HPV16 E6-E7 protein.** HeLa cells were transfected with mRNA encoding HPV16 E6-E7 antigens with MITD (E6E7-MITD), hCD1d (E6E7-hCD1d), KDEL (E6E7-KDEL) or no localization scaffold (E6E7) or control RNA (control). Expression of E6-E7 protein was measured by flow cytometry **(A)** and confocal microscopy **(B)**. **A.** For flow, surface **(left)** and intracellular **(right)** expression were measured using mouse anti-HPV16 E7 antibody and anti-mouse IgG secondary. **B.** By confocal, colocalization of HPV16 E6-E7 with subcellular compartments was assessed through co-staining HPV16 E6-E7 protein with early endosomes (anti-EEA1), lysosome (anti-LAMP1), Golgi complex (anti-Golgin97), endoplasmic reticulum (anti-SERCA2), and late endosomes (anti-VPS25) and analysis of Z-stack images over time. E6-E7 expression quantified as overlapped volume ratio for each construct by each subcellular compartment **(left panels)**. For each construct, mean fluorescence intensity of E6-E7 expression and for each organelle marker was compiled over time **(right panels)**.

**Figure S3: Immune activation data for individual donors.** HEK293 cells were transfected with mRNA encoding IL-12, LIGHT, or both (IL-12+LIGHT), or non-coding mRNA (Ctr). Purified CD3+ T cells from healthy donors (n=8; 202C, 302C, 801C, 802C, 201C, 010, 368, 2900) were incubated with anti-CD3 antibody and co-cultured with mRNA-transfected HEK293 cells. Proliferation of CD8 and CD4 T cells was measured by CTV dye dilution by flow after incubation for 4 days. Example of proliferation from single donor shown in **(A)**. Fold increase was calculated for each donor over proliferation following treatment with anti-CD3 only (CD3) **(B)**.

**Figure S4: Protection from tumor rechallenge is CD8+ T cell dependent.** mNTX250-treated mice that cleared primary C3.43 tumor challenge were administered anti-CD8 antibody (mNTX250 + aCD8; n=5) or isotype control (mNTX250 + isotype; n=5) three times on consecutive days, starting 89 days post initial tumor challenge. CD8+ T cell depletion in blood was assessed by flow cytometry staining for representative animals from mNTX250 + aCD8 (Mouse #1 and #2), mNTX250 + isotype (Mouse #3 and #4), and a naïve mouse **(A)**. Anti-CD8 and isotype control-treated mice (n=5) were re-implanted with C3.43 tumors subcutaneously on the opposite flank. Naïve C57BL/6 mice (n=5) were implanted as controls. Tumor growth measured over time **(B)**. Error bars represent the mean ± SEM.

### *Supplemental Information: Generation of membrane stabilized Engineered LIGHT (ENG-LIGHT).*

Based on the observed contribution of LIGHT as a T-cell co-stimulator, we decided to explore a version of LIGHT that is membrane-anchored to potentially enhance its immunomodulatory effects. TNFSF14 (LIGHT) is a type II transmembrane protein of the TNF superfamily that exists in both membrane-bound and soluble forms. The membrane-bound form is a homotrimer expressed on activated lymphocytes and other immune cells, while the soluble form arises via proteolytic cleavage of the extracellular domain by metalloproteinases. This shedding mechanism is like that seen in other TNF superfamily members such as TNFα and FasL and is mediated through a cleavage site encoded in exon 2. LIGHT exerts immunomodulatory functions primarily through cis- or trans-interactions with HVEM or lymphotoxin beta receptor (LTβR), respectively. Importantly, LIGHT's co-stimulatory activity on T cells requires displacement of the inhibitory receptor BTLA from HVEM. Previous studies indicate that membrane-bound LIGHT, but not soluble LIGHT, can effectively displace BTLA, thereby enhancing T cell activation and proliferation[28–31]. Indeed, immobilized LIGHT induces stronger T cell responses than its soluble counterpart (Supplementary Figure 5A). To augment these properties, we hypothesized that a membrane-stabilized form of LIGHT resistant to proteolytic cleavage would exhibit superior activity. We engineered multiple LIGHT variants by replacing regions within the protease-sensitive site with either flexible or rigid linkers. Through functional screening, we identified a well-expressed and functionally active, membrane-stabilized variant—ENG-LIGHT—for further analysis.

### *ENG-LIGHT Enhances Membrane Retention and Reduces Soluble Shedding*

HEK293 cells were transfected with mRNA encoding either wild-type (WT) LIGHT or ENG-LIGHT. Membrane-bound and soluble LIGHT were quantified in cell lysates and supernatants, respectively. ENG-LIGHT exhibited increased surface expression and significantly reduced soluble shedding compared to WT LIGHT (Supplementary Figure 5B).

### *Soluble LIGHT Diminishes HVEM Signaling by Competitive Inhibition*

To determine the functional consequence of soluble LIGHT on HVEM signaling, HEK293 cells transfected with WT LIGHT were co-cultured with HVEM reporter cells in the presence of increasing concentrations of recombinant soluble LIGHT. The reporter cells express luciferase under an NF-κB promoter. Increasing soluble LIGHT levels reduced HVEM activation by competing with membrane-bound LIGHT for receptor binding, leading to reduced luciferase activity (Supplementary Figure 5C).

### *ENG-LIGHT Elicits Enhanced HVEM Signaling and Pro-Inflammatory Cytokine Production*

To directly compare HVEM activation by WT and ENG-LIGHT, HEK293 cells transfected with the same amount of mRNA encoding each variant were co-cultured with HVEM reporter cells. ENG-LIGHT-expressing cells induced significantly stronger HVEM-mediated luciferase expression than WT LIGHT (Supplementary Figure 5D). A murine version of ENG-LIGHT (mENG-LIGHT) was then tested in vivo. Mice bearing subcutaneous MC38 tumors received two doses of IL-12 mRNA in combination with either mENG-LIGHT or WT LIGHT mRNA three days apart. Tumors were harvested 12 hours after the second dose for cytokine analysis. mENG-LIGHT-treated tumors exhibited significantly higher levels of IFNγ, TNFα, and IL-15, normalized to total protein content, compared to WT LIGHT-treated tumors (Supplementary Figure 5E).

Notably, we engineered a membrane-stabilized variant of LIGHT (ENG-LIGHT) that prevents proteolytic shedding, thereby maintaining its functional membrane-bound state and increasing HVEM-mediated signaling. In both in vitro and in vivo assays, ENG-LIGHT induced stronger immune activation and cytokine production than wild-type LIGHT, underscoring its utility in maintaining localized immune activation while minimizing systemic toxicity. Compared to previous approaches that used soluble or wild-type forms of TNFSF ligands[22], the final version of NTX250 incorporates a genetically stabilized LIGHT variant, providing superior control of signaling dynamics and tissue specificity.

**Figure S5: Membrane-stabilization of mRNA-encoded LIGHT enhances costimulatory function. A.** T cells isolated from healthy donor PBMCs were labeled and then incubated in 96-well plates pre-treated with anti-CD3 (CD3) and/or LIGHT protein at indicated concentrations. Soluble LIGHT (sbl) or agonistic anti-CD28 (CD28) were included as controls. Next day, proliferation **(left panels)** and CD25 upregulation **(right panels)** of CD8+ **(top panels)** and CD4+ **(lower panels)** T cells were measured by flow cytometry. **B, C, D.** HEK293 cells were transfected with mRNA encoding wildtype (wt LIGHT) or engineered (ENG LIGHT) LIGHT or untreated. Levels of soluble LIGHT were measured in transfection supernatants by ELISA **(B, left panel)**. Expression of membrane-associated LIGHT on cell surface was measured by flow cytometry **(B, right panel)**. Jurkat HVEM reporter cells were incubated with increasing amounts of soluble LIGHT protein prior to co-incubation with ENG-LIGHT mRNA transfected HEK293 cells **(C)**. ENG-LIGHT mRNA and wt LIGHT mRNA transfected HEK293 cells **(D)** were incubated with Jurkat HVEM reporter cell line and HVEM activation measured by luciferase activity (RLU). **E.** C57BL/6 mice (n=5) were implanted with MC38 tumors subcutaneously. Nutshell™ formulations of mRNA encoding mIL-12 alone (IL-12 mRNA) or in combination with wt mLIGHT (IL-12+wt LIGHT) or with membrane-stabilized mLIGHT (IL-12+ENG-LIGHT) or control mRNA (Ctr mRNA) were administered intratumorally on D0 and D3. 12h post-2nd dose, tumors were removed and tumor supernatant collected. IFNγ, TNFα, and IL-15 were measured in tumor supernatant by Luminex assay. Statistical significance was determined by two-tailed, Student's t-test: *$^{*}p<0.05$, $^{**}p<0.01$, $^{***}p<0.001$, $^{****}p<0.0001$*
